# Age- and Sex-Dependent Caveolar Gene Expression in Normoxic and Post-Ischemic Mouse Hearts

**DOI:** 10.1101/2024.03.01.582960

**Authors:** Can J. Kiessling

## Abstract

Age and sex modify myocardial I-R tolerance with caveolar invaginations in the plasma membrane shown to be important determinants of ischemic tolerance as well as responsiveness to cardioprotective stimuli. Caveolar membrane domains are formed by variety of coat proteins including those of caveolins, cavins and Popeye domain-containing family of proteins. Expression of these proteins are unclear in the aging male and female myocardium. Expression of caveolae-associated transcripts and proteins were assayed in normoxic and post-ischemic (20 min ischemia/60 min reperfusion) myocardium from 8-, 16-, 32- and 48-week old male and female mice (C57Bl/6). Using cultured HL-1 cells, the reliance of acute A_1_ Adenosine Receptor agonism on caveolae was also assessed, a known stress tolerance determinant that reside in caveolae. Significant down-regulation of chief caveolae transcripts *Cav3* and *Cavin1* correlated with 38%-52% reductions in Caveolin-3 & Cavin-1 protein in normoxic middle-aged hearts (respectively), congruent with age-related repression of caveolins in other tissue types. For females, caveolae-related genes were largely stable with aging which were consistent with previous findings that middle-aged female hearts display increase stress tolerance as measured by LDH release following I-R. Age thus appears to suppress transcription/expression of caveolar proteins critical to myocardial responses to stress and protective stimuli, an effect more pronounced in male vs. female myocardium. These age-dependent shifts in caveolar coat protein expression was associated with exaggerated contractile dysfunction associated evident at 32- and 48-weeks (vs. 8 weeks) in male hearts, but not associated with impaired recovery in 32 and 48-weeks (vs. 8 and 16 weeks) in females. In HL-1 cardiomyocytes subject to simulated ischemia-reperfusion, acute A_1_AR agonism with 100 nM 2-chloro-N6-cyclopentyadenosine (CCPA) resulted in cytoprotection evident by significantly decreased LDH release (by ̴20%), which was abolished by caveolae & cholesterol depletion agent methyl-β-cyclodextrin (MβCD, 1 mM), suggesting caveolar involvement in protective signaling.

**Highlights:**

- Aged-related reduction in ischemic tolerance evident in middle-aged male and female heart before “aged” phenotype typically used in aging studies.
- Declining ischemic tolerance coincident with age-related shifts in *Cav1*, *Cav3*, *Cavin1* and *Popdc1* mRNAs, and reduced Caveolin-3 and Cavin-1 protein in males.
- In the aging female myocardium crucial caveolar coat transcripts *Cav3, Cavin1 and Popdc1* were largely stable with aging and I-R, consistent with reports of enhanced stress tolerance in the female heart & role of caveolae in promoting stress resistance.
- Top predicted microRNA regulators of *Cav3* transcript microRNA-22 and microRNA-101b did not inversely correlate with observed *Cav3* mRNA expression profiles in the aging heart.
- In simulated ischemia-reperfused cultured cardiomyocytes, acute A_1_AR agonism markedly decreased cell injury, this protection was abolish by cholesterol & caveolae depleting agent MβCD which also enhanced cell death, highlighting the role of caveolae in endogenous ischemic tolerance.

## Introduction

Aging is an ill-defined and multifaceted process that results in complex molecular and physiological changes. This is particularly relevant in the context of heart disease, which is highly age-dependent: age strongly influences the response of the heart to both disease/insult (Lakatta, 2015) and to potential protective therapies (Boengler et al., 2008; Peart et al., 2014; Randhawa et al., 2018). Sex is also a critical factor, with epidemiological and experimental evidence that hearts from pre-menopausal females are resistant to ischemia-reperfusion (I-R) injury compared with hearts from age-matched males (Korzick and Lancaster, 2013; Willems et al., 2005). Specialized caveolar microdomains within the plasma membrane are critical to the hearts ability to withstand injury during I-R, and to respond to protective stimuli (Tsutsumi et al., 2008; See Hoe et al., 2014; Horikawa et al., 2008). Caveolae compartmentalize and concentrate various classes of signaling molecules, including determinants of cellular survival, death, mechanoprotection, remodeling and substrate metabolism (Parton, 2018; Das, Cui and Das, 2007). They localize receptor tyrosine kinases, G protein-coupled receptors (GPCRs) and their molecular effectors (Insel et al., 2005) and play roles in regulating substrate metabolism and cellular responses to mechanical loading/stretch (Parton and del Pozo, 2013; Sinha et al., 2011).

The chief structural proteins of caveolae are the caveolins, consisting of three members: caveolin-1 (Cav-1), caveolin-2 (Cav-2) and caveolin-3 (Cav-3), encoded by *Cav1*, *Cav2* and *Cav3* genes, respectively (Williams and Lisanti, 2004). Although the heart expresses all 3, cardiomyocyte caveolae are specifically dependent on expression of muscle-specific Cav-3, with Cav-3-deficiency or -overexpression ablating or increasing myocardial caveolae formation, respectively (Park et al., 2002; Tsutsumi et al., 2008). Pharmacological disruption (via Methyl-β-cyclodextrin, MβCD) of caveolae similarly impairs ischemic tolerance, while *Cav3* gene overexpression and knockout models confirm that cardioprotection via preconditioning and other stimuli is reliant on caveolar microdomains (Tsutsumi et al., 2008; See Hoe et al., 2014; Horikawa et al., 2008) while nitration of Cav-3 at Tyr73 has been shown to result in signal complex dissociation leading to cardiac insulin/adiponectin resistance in the prediabetic heart, subsequently contributing to ischemic heart failure progression (Meng et al., 2023). Age-related changes in membrane cholesterol (particularly enriched in caveolae) and Cav-3 protein are present in 74-week old (“aged”) myocardium, and linked to stress-intolerance and refractoriness to protective stimuli (Peart et al., 2014). Recent RNA-seq studies revealed that cardiac transcriptomics are significantly different across the life course for 1788 transcripts in juvenile vs. aged hearts (Yusifov et al., 2021). Differences in caveolar protein levels and function may also underlie sex-dependent stress responses, with evidence increased Cav-3 association of protective eNOS may contribute to the cardioprotective phenotype in female vs. male hearts (Sun et al., 2006).

More recently additional coat proteins have been identified, including Cavins 1-4 and Popeye domain-containing proteins (Popdc1, Popdc2, Popdc3), which contribute to caveolae formation and morphology in a tissue-specific manner (Bastiani et al., 2009; Alcalay et al., 2013; Hansen et al., 2013). Thus far only Cavin-1 (initially identified as polymerase I and transcript release factor - PTRF) and Popdc1 (also known as blood vessel epicardial substance - BVES) have been shown to influence cardiomyocyte caveolae formation. Cavin-1 has also been shown to influence cell death processes (McMahon et al., 2019). Similar to Cav-3, deficiencies in Cavin-1 or Popdc1 reduce caveolar density and are associated with impaired stress-tolerance and reduced efficacy of protective stimuli (Alcalay et al., 2013; Taniguchi et al., 2016; Kaakinen et al., 2017). While age-related changes in caveolins have been documented in several tissues, concurrent effects of age and sex on myocardial caveolae-associated protein expression profiles are yet to be detailed (Lowalekar et al., 2012; Barrientos et al., 2015; Head et al., 2010). We therefore investigated the age and sex dependencies of caveolin, cavin and Popdc transcription and expression in normoxic and post-ischemic mouse hearts. As Caveolin-3 transcript was found to be consistently down-regulated with aging as early as 16-weeks old in the male hearts, microRNAs (miRNAs/miRs) predicted to target Cav-3 gene were assessed in the aging myocardium. Previously A_1_ Adenosine Receptor (A_1_AR) activation has been shown to result in cytoprotective signaling in HL-1 cells (Williams-Pritchard et al., 2011), although it remained to be shown whether caveolar depletion could abolish A_1_AR-mediated cytoprotection. Hence, the reliance of A_1_AR on caveolar microdomains for cytoprotective signaling was investigated as A_1_AR associate with caveolae and caveolin-3 in ventricular cardiomyocytes (Lasley, 2010).

## Materials & Methods

### Perfused mouse heart model

Hearts were obtained from male and female C57Bl/6 mice aged 8, 16, 32 and 48 weeks (*n*=8-12/group). Collection of hearts was approved by and performed in accordance with the guidelines of the Animal Ethics Committee of Griffith University, which is accredited by the Queensland Government, Department of Primary Industries and Fisheries under the guidelines of “The Animal Care and Protection Act 2001, Section 757”. Mice were anesthetized with 60 mg/kg sodium pentobarbital, hearts excised into ice-cold Krebs-Henseleit solution and either trimmed to ventricular myocardium and frozen at -80°C for later analysis, or Langendorff perfused to assess responses to I-R, as detailed previously (Willems et al., 2005; Headrick et al., 2003; Ashton et al., 2023). Hearts were subjected to 20 minutes of global normothermic ischemia followed by 60 minutes aerobic reperfusion. Coronary venous effluent was sampled throughout the post-ischemic period, stored at 4°C and assayed the following day for lactate dehydrogenase (LDH) activity as a measure of cell death using a Cytotox 96® Non-radioactive Cytotoxicity Assay (Promega, Madison, Wisconsin, USA).

### RNA isolation from left ventricular tissue

Atrial and vascular tissue was removed and left ventricular myocardium dissected from each heart and homogenized in QIAzol™ reagent (QIAGEN, Maryland, USA). Total RNA containing microRNA was isolated according to manufacturer’s guidelines using miRNeasy spin columns (QIAGEN, Maryland, USA). The RNA yield and purity were determined using a NanoDrop ND-1000 (Thermo-Fisher, Massachusetts, USA). RNA integrity was assessed using a 2100 Bioanalyzer (Agilent Technologies, Santa Clara, CA, USA), with RNA integrity (RIN) scores ≥8.0 for each sample. Total RNA was stored at -80°C prior to cDNA synthesis. For cDNA synthesis 500 ng of total RNA was reverse transcribed using the Superscript III First Strand cDNA Synthesis System (Life Technologies, Carlsbad, CA, USA) according to manufacturer’s protocols. Synthesized cDNA was diluted with nuclease free water 1:20 and stored at -20°C before use.

### RT-qPCR of Caveolin, Cavin and Popdc

Two-step RT-qPCR was performed on a CFX96 Real-Time PCR Detection System (Bio-Rad, Hercules, CA) utilizing SYBR Green I chemistry. PCR primers (see Supplementary Table 1) were designed with PerlPrimer software (Marshall, 2004) according to standard guidelines (Ashton and Headrick, 2007). The final reaction volume (10 μL) included 1X iQ SYBR Green Supermix (Bio-Rad, Hercules, CA), 100 nM of each primer and 5 μL of diluted (1:20) cDNA. Optimal qPCR cycling conditions consisted of an initial denaturation at 95°C for 3 minutes followed by 40 cycles of 95°C for 15 seconds and 61°C for 60 seconds. After the final PCR cycle, reactions underwent melt-curve analysis to detect non-specific amplicons. Resultant amplicons were checked for predicted amplicon sizes using a QIAxcel electrophoresis system (QIAGEN, Maryland, USA). All reactions were performed in triplicate with each plate containing an equal number of samples from each group (N=6), a calibrator control derived from a pool of all cDNA samples, and a no template control. PCR amplification efficiencies for each primer pair were calculated using a 5-log serial dilution of the calibrator sample. PCR data were analyzed using CFX Manager v2.0 (Bio-Rad, Hercules, CA). Following automatic baseline correction, the threshold level was set during the geometric phase of PCR amplification for quantitative measurement. The calibrator sample was used to normalize inter-assay variation, with threshold coefficients of variance for intra- and inter-assay replicates <1% and <5%, respectively. Normalized expression (ΔΔCq) was calculated, with expression normalized to reference gene levels and log_2_ transformed and expressed relative to 8 week-old values.

For reference gene selection five additional genes (*Actb*, *Gapdh*, *Pgk1*, *Ppi* and *Rpl13a*) were assessed using Bio-Rad CFX Manager Target Stability analysis to determine their usability as reference genes in the aging male and female hearts. Of the reference genes assessed for stability, there was no single reference gene that was stably expressed with age, I-R and sex. *Pgk1* was chosen as a normalizer as it was shown to exhibit the most stable expression with age and I-R in both male and female hearts thus permitting within-sex normoxic vs. I-R comparisons. We noted that the commonly used reference gene *Gapdh* was unstable (M-value > 1.0) in the post-ischemic female hearts (data now shown, available on request).

### In-silico bioinformatics & RT-qPCR of Cav3-related microRNAs

For predicted microRNAs targeting the *Cav3* transcript, in-silico prediction using TargetScan (mouse, https://www.targetscan.org/vert_80/) was used to screen for two top candidates of *Cav3* supplemented with literature search of these microRNAs for evidence of gene expression regulation. Mature microRNA sequences were obtained from the miRBase database (https://mirbase.org/) and used as the template to design the microRNA specific primer (Supplement Table 1). OligoAnalyzer (https://www.idtdna.com/pages/tools/oligoanalyzer) (IDT, Iowa, USA) was used to design the microRNA primers at Tm of 58*C with BLAST used to ensure primer specificity to the microRNA target. Following microRNA extraction using miRNeasy columns (QIAGEN, Maryland, USA), the NCode miRNA First-Strand cDNA Synthesis Kit (Thermo-Fisher, Massachusetts, USA) was used to synthesize cDNA from 1.0 µg of cardiac total RNA containing the microRNA. Due to their small size (22nt) the mature microRNA first requires polyadenylation, secondly an oligo-dT adapter primer is used to prime the cDNA synthesis reaction. This adapter primer contains a unique proprietary sequence at its 5’ end, which allows for amplification of cDNA using a complimentary universal primer paired with a primer specific to the microRNA sequence of interest. Forward primers used in this experiment to amplify resultant cDNA containing microRNA libraries are shown in supplement table 1. For microRNA expression analysis, optimal qPCR cycling conditions consisted of an initial denaturation at 95°C for 3 minutes followed by 40 cycles of 95°C for 15 seconds and 58°C for 30 seconds. After the final PCR cycle, reactions underwent melt curve analysis to detect non-specific amplicons. Reference gene selection for gene expression normalization was performed using CFX-Manager (Bio-Rad, Hercules, CA) geNorm analysis. The reference panel was composed of *miR-24*, *miR-26*, *miR-103*, *miR-191*, *Rnu1a1* and *Rnu6*, with *miR-191* shown to be most stable in expression across both the normoxic and the post-ischemic hearts (data not shown, available on request).

### Western immunoblotting

Immunoblot analysis was employed to confirm myocardial expression of select transcript products. Antibody for β-actin (#4967S, rabbit polyclonal) was obtained from Cell Signaling (Danvers, MA, USA), Caveolin-3 (sc-5310, mouse monoclonal) from Santa Cruz Biotechnology (Dallas, TX, USA), and Cavin-1 (ab48824, rabbit polyclonal) from Abcam (Cambridge, MA, USA). Samples were prepared in fresh Laemmli buffer containing 30 μg of protein from whole-cell fractions and loaded onto 12% acrylamide gels, with equal loading confirmed by Ponceau staining and β-actin expression. Electrophoresis was carried out at 75V for 90 minutes. Protein was transferred to an Immobilon®-FL PVDF membrane (Merck Millipore, Billerica, MA, USA) and blocked in Licor blocking buffer for 120 minutes (Licor, Lincoln, NE, USA). Membranes were then incubated with primary antibody (Caveolin-3, 1:1500; Cavin-1, 1:2000; β-actin: 1:1500) overnight at 4°C. Dual colored labelling was performed using fluorescently labelled secondary antibodies. Briefly, membranes were incubated in a 1:20,000 dilution of IRDye® 680RD donkey anti-rabbit IgG and IRDye® 800CW goat anti-mouse IgG (Licor, Lincoln, NE, USA) secondary antibodies for 120 min with gentle agitation. The labeled membranes were washed with TBS and TBST and scanned using an ODYSSEY® CLx infrared scanner (Licor, Lincoln, NE, USA). Densitometry analysis was performed using Image Studio Lite™ 5.2 (Licor, Lincoln, NE, USA), with Caveolin-3 and Cavin-1 expression normalized to stably expressed loading control (β-actin). Expression ratios were expressed relative to 8 week-old values.

### Cell Culture & Simulated Ischemia in HL-1 cells

Cells of the murine atrial-derived cardiac cell line HL-1 were a kind gift from Professor William C. Claycomb (Louisiana State University, Baton Rouge, LA). Cells were grown in culture vessels pre-coated with 0.00125% fibronectin in 0.02% gelatin and maintained in Claycomb medium (Sigma Aldrich, St. Louise, MO, USA) supplemented with 10% fetal bovine serum (Life Technologies, Carlsbad, CA, USA), 0.1 mM norepinephrine, 2 mM L-glutamine, 100 U/mL penicillin/streptomycin, and 0.25 µg/mL amphotericin B. Cells were cultured at 37°C in a humidified atmosphere of 5% CO2 and 95% air. Media was replaced every second day and cells passaged at >80% confluency when required using 0.25% Trypsin-EDTA (Life Technologies, Carlsbad, CA, USA). Cells were screened every two months for mycoplasma contamination using the MycoAlert™ Mycoplasma Detection Kit (Lonza, Basel, Switzerland). Passaged cells were continually monitored for viability using Countess cell counting slides (Thermo-Fisher, Massachusetts, USA). Simulated I-R in cultured HL-1 myocytes was performed similar to a previous report (See Hoe et al. 2014) with 3 hour of ischemia followed by 5 hour of reperfusion. Sealed hypoxic chamber (STEMCELL Technologies, Vancouver, BC, Canada) and the oxygen monitor (Maxtec Inc, Salt Lake City, Utah) used in HL-1 cardiomyocyte simulated I-R experiments are shown below. Cells were untreated or treated with 100 nM 2-chloro-N6-cyclopentyadenosine (CCPA) (Sigma-Aldrich, St. Louis, Missouri, United States) for 30 min for CCPA only group, 1 mM methyl-β-cyclodextrin (MβCD) (Sigma-Aldrich, St. Louis, Missouri, United States) for 60 min for MβCD treatment only group, and 30 minutes of 1 mM MβCD prior to 30 min incubation with 100 nM CCPA for MβCD + CCPA treatment group before simulated I-R. LDH activity was assayed using a CytoTox 96 assay kit (Promega, Madison, Wisconsin, USA) according to the manufacturer’s instructions. Normoxic control cells incubated with MβCD showed no detectable cytotoxicity based on LDH changes and flow cytometric analysis of 7-AAD/PI and Annexin V-PE staining (data not shown).

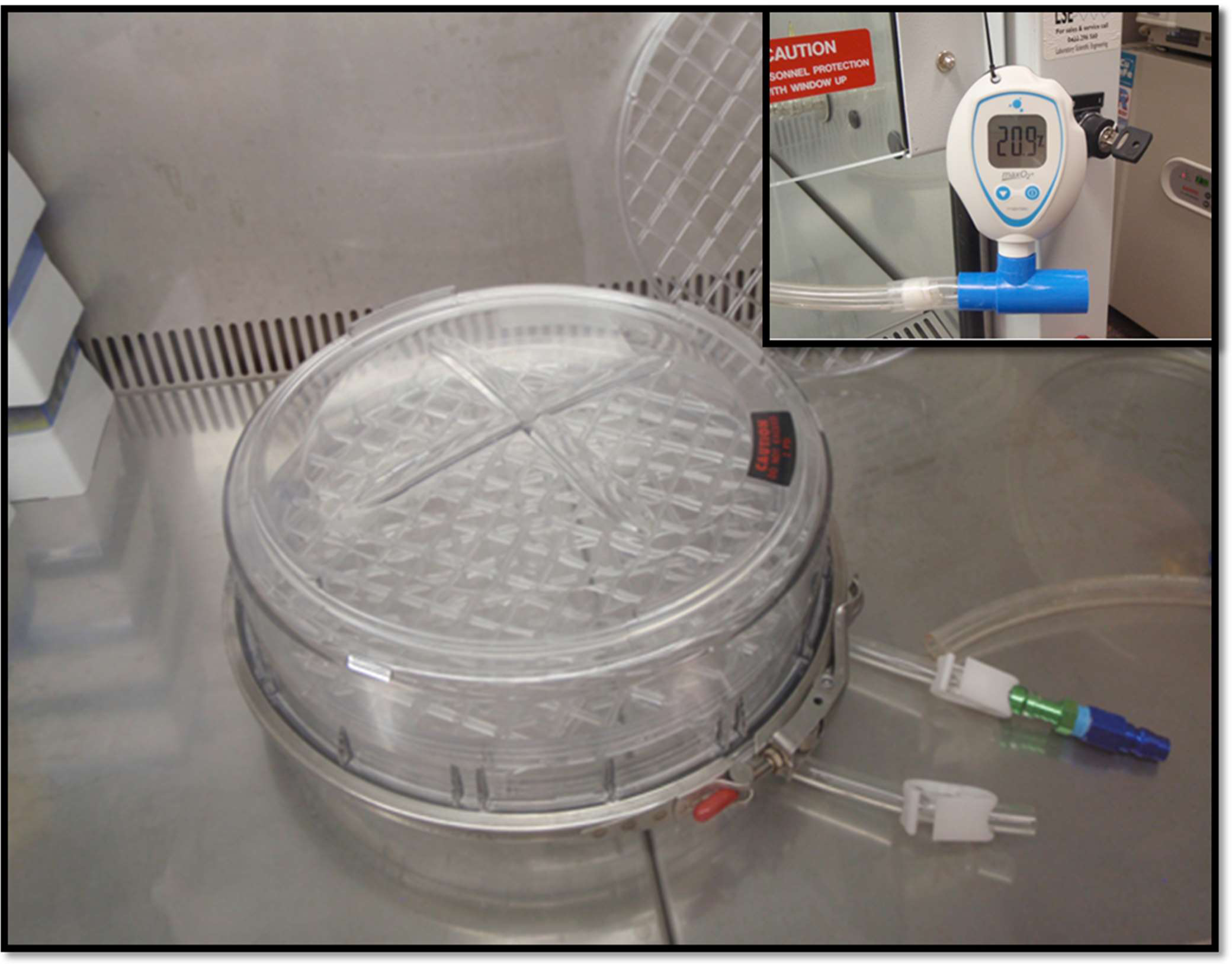

### Statistical analyses

LVDP recovery, LDH release and immunoblot analysis are expressed as means ± the standard error of the mean (SEM). Statistical analyses were performed and plotted using Prism 10.2 software (GraphPad, California, USA). For pair-wise comparisons in protein expression data a two-tailed Student’s t-test was employed. Parametric data comparisons between more than two groups were made via an analysis of variance (ANOVA), with Tukey’s post-hoc test applied where differences were detected in gene expression analysis, LDH release and post-ischemic recovery. For age-matched comparisons (normoxia vs. I-R), Sidak’s post-hoc test was applied where differences were detected in gene expression. When standard deviation differed significantly Brown-Forsythe and Welch ANOVA was used to detect for significance with Dunnett T3 for post-hoc detection of differences. Statistical significance was accepted for *P*<0.05.

## Results

### Effects of age on post-ischemic cardiac dysfunction and injury in male and female hearts

Baseline (normoxic) contractile function did not differ significantly across age groups, or in males vs. females (Table 1). Consistent with previous observations (Peart et al., 2014; Willems et al., 2005; Ashton et al., 2023), flow rate declined with age in male and female hearts at 32 and 48-weeks (vs. sex-matched 8-week) (Table 1). Functional resistance to I-R was assessed from recoveries of left ventricular pressure development. Following 20 min ischemia/60 min reperfusion, significant age-related reductions in functional recovery were evident in female hearts whereas male hearts showed age-related declines that that were not statistically significant (Figure 1A). Specifically, contractile dysfunction was exaggerated in 32- and 48-week-old male mice (vs. 8-week), while in female myocardium dysfunction was significantly greater in 48- vs. 8- and 16-week hearts (Figure 1A). Myocardial I-R injury assessed from Lactic Acid Dehydrogenase (LDH) efflux was markedly exaggerated in middle-aged hearts for both male (48 vs. 8) and female (48 vs. 8 and 16) mice (Figure 1B).

**Figure 1.**
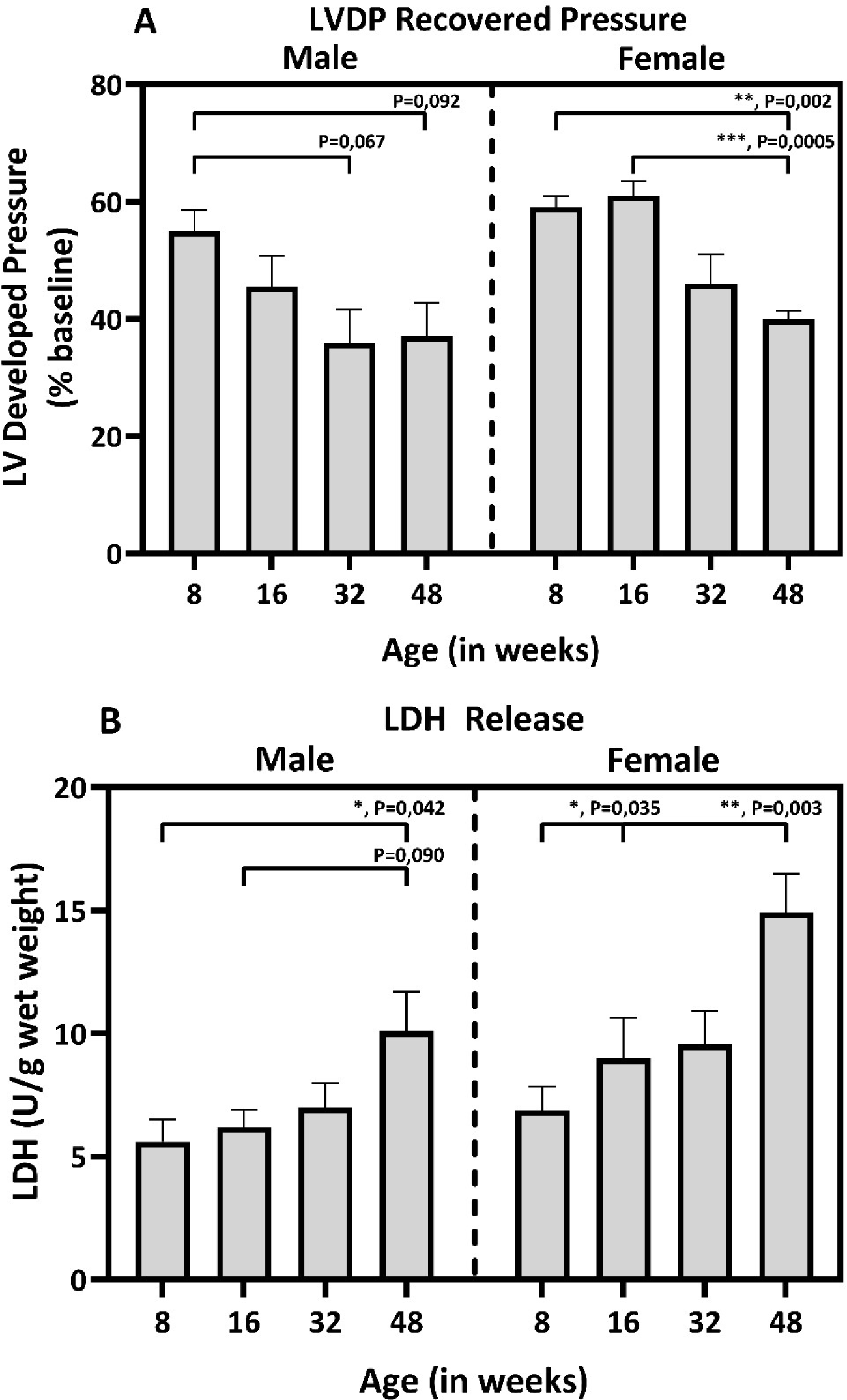
Functional recovery and cell death following 20 min ischemia and 60 min reperfusion in hearts from young to middle-aged male and female mice. Bar graphs present: **A)** recovery of left ventricular developed pressure; and **B)** post-ischemic efflux of LDH for hearts from male (left) and female (right) mice (8, 16, 32 and 48 weeks of age) (*n*=6/group). Data are expressed as mean ± SEM. *, *P*<0.05, **, *P*<0.01, ***, *P*<0.001.

**Table 1.** Baseline (pre-ischemic) functional parameters for hearts from 8, 16, 32, 48-week old male and female mice.

| **Sex** | **Age**<br>**(weeks)** | **EDP**<br>**(mmHg)** | **LVDP**<br>**(mmHg)** | **Intrinsic Rate**<br>**(bpm)** | **+dP/dt**<br>**(mmHg/s)** | **-dP/dt**<br>**(mmHg/s)** | **Flow**<br>**(ml/min/g)** |
| --- | --- | --- | --- | --- | --- | --- | --- |
| Male | 8 | 4±2 | 158±11 | 331±12 | 6369±281 | -4092±160 | 28.9±1.5 |
|  | 16 | 3±2 | 150±10 | 317±13 | 6598±460 | -4368±164 | 26.7±2.3 |
|  | 32 | 5±1 | 153±5 | 318±7 | 6001±489 | -4479±247 | 19.4±2.3* |
|  | 48 | 4±2 | 158±5 | 311±9 | 6623±386 | -3921±217 | 22.8±2.2* |
| Female | 8 | 3±1 | 151±8 | 344±13 | 7030±417 | -4698±239 | 29.6±1.8 |
|  | 16 | 3±2 | 158±7 | 331±12 | 6950±349 | -4513±240 | 28.7±2.5 |
|  | 32 | 4±1 | 153±8 | 323±10 | 6324±401 | -3921±237 | 22.0±1.8* |
|  | 48 | 4±2 | 155±9 | 330±9 | 5685±397 | -3755±306 | 22.0±1.9* |

Data acquired after normoxic stabilization ( $n=6$ per group). EDP, end-diastolic pressure; LVDP, left ventricular developed pressure; Intrinsic Rate, intrinsic heart rate prior to ventricular pacing (at 420 bpm); dP/dt, differential of left ventricular pressure development or relaxation over time; flow, coronary flow rate. Data are expressed as mean ± SEM ( $n=6$ ). \*, $P<0.05$ vs. 8 week sex-matched.

### Age-dependence of caveolar protein transcription & predicted microRNA regulators of *Cav3*

Prior to gene expression analysis of caveolins, cavins and Popdcs with cardiac aging, cell-specific gene expression patterns for caveolins, cavins and Popdcs were screened in left ventricular homogenates, 3T3 fibroblasts and cultured atrial HL-1 cardiomyocytes. Left ventricular isolates showed relatively high expression of all members of caveolins, cavins and Popdcs (supplementary Figure S1). Concordant with literature, muscle-specific expression *Cav3 and Popdcs (1-3)* were confirmed in left ventricular samples and atrial HL-1 cardiomyocytes with 3T3 fibroblasts showing non-detectable expression. Interestingly, *Popdc1* & *Popdc2* were expressed in both HL-1 atrial cardiomyocytes and left ventricular samples whereas *Popdc3* expression was exclusive in left ventricular isolates (supplementary Figure S1). *Cavin4* (aka **Mu**scle-**R**estricted **C**oiled-Coil Protein/MURC) displayed low-level expression in 3T3 fibroblast even though tissue analysis (via https://www.proteinatlas.org/ENSG00000170681-CAVIN4) suggested skeletal and cardiac-type specificity (Bastiani et al., 2009; Hansen et al., 2013).

#### Caveolin transcripts

Two-way ANOVA revealed a statistically significant main effect of age on expression of both *Cav1* (*F*(3,40) = 5.56, *P*=0.003) and *Cav2* (*F*(3,40) = 4.69, *P*=0.007) in male hearts. These changes occurred exclusively in normoxic hearts, with aging reducing expression of *Cav1* by 1.8-fold (48 vs. 8-week) and *Cav2* by 1.4-fold (48 vs. 8-week) (Figure 2). No age-related differences in *Cav1* and *Cav2* expression were observed in post-ischemic male hearts. However, for *Cav3* expression, a statistically significant interaction between the effects of age and ischemia was observed (*F*(3,40) = 11.01, *P*<0.0001). More specifically, significant down-regulation of *Cav3* was observed in the normoxic heart at 16-weeks compared to 8-weeks old weeks (1.5-fold). This repression was maintained in older hearts (Figure 2). In post-ischemic male hearts this down-regulation was not as pronounced, however a 1.5-fold down-regulation was observed in the 48-week old vs. 16-week old hearts further supported by the significant effect for I-R identified by two-way ANOVA analysis (*F*(1,40) = 32.87, P<0.0001). In addition, in the post-ischemic heart *Cav3* expression is induced above baseline (1.5-fold). As the heart ages this response appears to be lost and an age-related decline in *Cav3* expression is observed in a similar manner to their normoxic counterparts.

**Figure 2.**
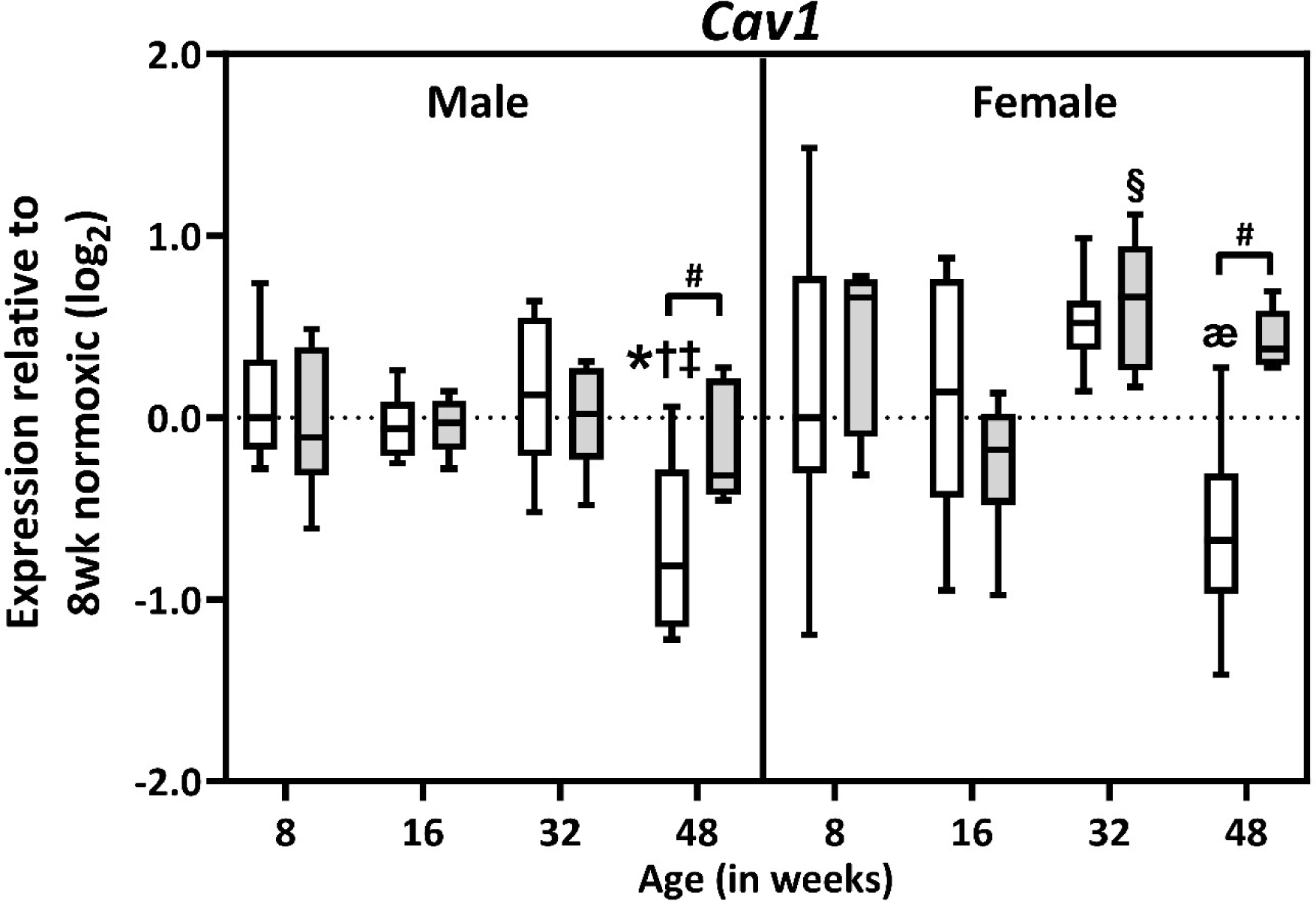
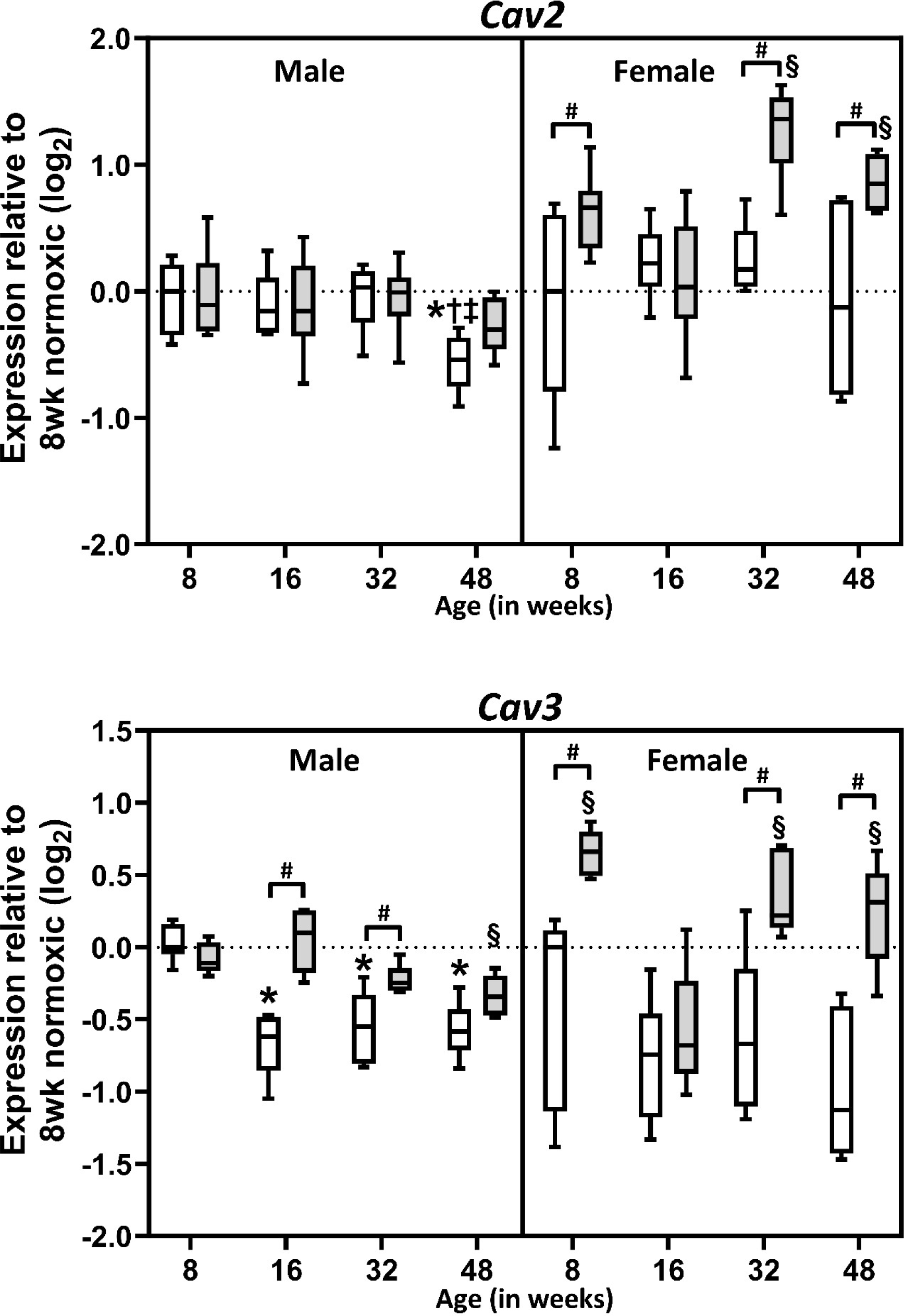
Age-dependent expression of caveolin transcripts in normoxic and post-ischemic myocardium. Shown are boxplots detailing expression changes for *Cav1, Cav2* and *Cav3* transcripts in normoxic (white) and post-ischemic (grey) male and female hearts (*n*=5-6/group). Boxes indicate the interquartile range (25%-75%) with the horizontal bar within each box indicating the median. Whiskers show minimum and maximum values. For normoxic hearts: *, *P* <0.05 vs. 8-week; †, P<0.05 vs. 16-week; ‡, P<0.05 vs. 32-week. For post-ischemic hearts: §, P<0.05 vs. 16-week; æ, P<0.05 vs. 32-week; #, P<0.05 vs. age-matched post-ischemic group (indicated by brackets).

The expression of caveolins was relatively independent of age in female (normoxic and post-ischemic) hearts (Figure 2), with a modest reduction in *Cav3* (significant effect of age; *F*(3,38) = 6.91, *P*=0.0008) not achieving statistical significance in post-hoc analysis (with a larger range of values in females). Induction of caveolin mRNA following I-R was also evident in females, although patterns of age-dependent expression of *Cav2* and *Cav3* differed (Figure 2).

#### Cavin transcripts

Age-related down-regulation (1.6-fold) of *Cavin1* was apparent in normoxic and post-ischemic hearts from male mice (48 vs. 8-week) (Figure 3), with a significant interaction of age identified (F(3,40) = 11.03, P=<0.0001). Post-ischemic *Cavin2* transcript also demonstrated a small (1.2-fold) yet significant decline in 48 vs. 8-week old post-ischemic hearts (Figure 3) (interaction with ischemia; F(1,40) = 35.30, P=<0.0001), and normoxic levels were up-regulated in 32 vs. 8-week hearts. Notably, *Cavin2* also demonstrated repression in age-matched post-ischemic vs. normoxic hearts (Figure 3) (16, 32 and 48-week), with a significant interaction between age and ischemia identified via 2-way ANOVA (F(3,40) = 6.02, P=0.0017). Expression of *Cavin3* and *Cavin4* transcripts were largely age-independent in normoxic and post-ischemic male hearts. Expression of cavin mRNA was also largely stable across ages in normoxic and post-ischemic female hearts, with an insignificant trend to *Cavin3* and *Cavin4* up-regulation (Figure 3).

**Figure 3.**
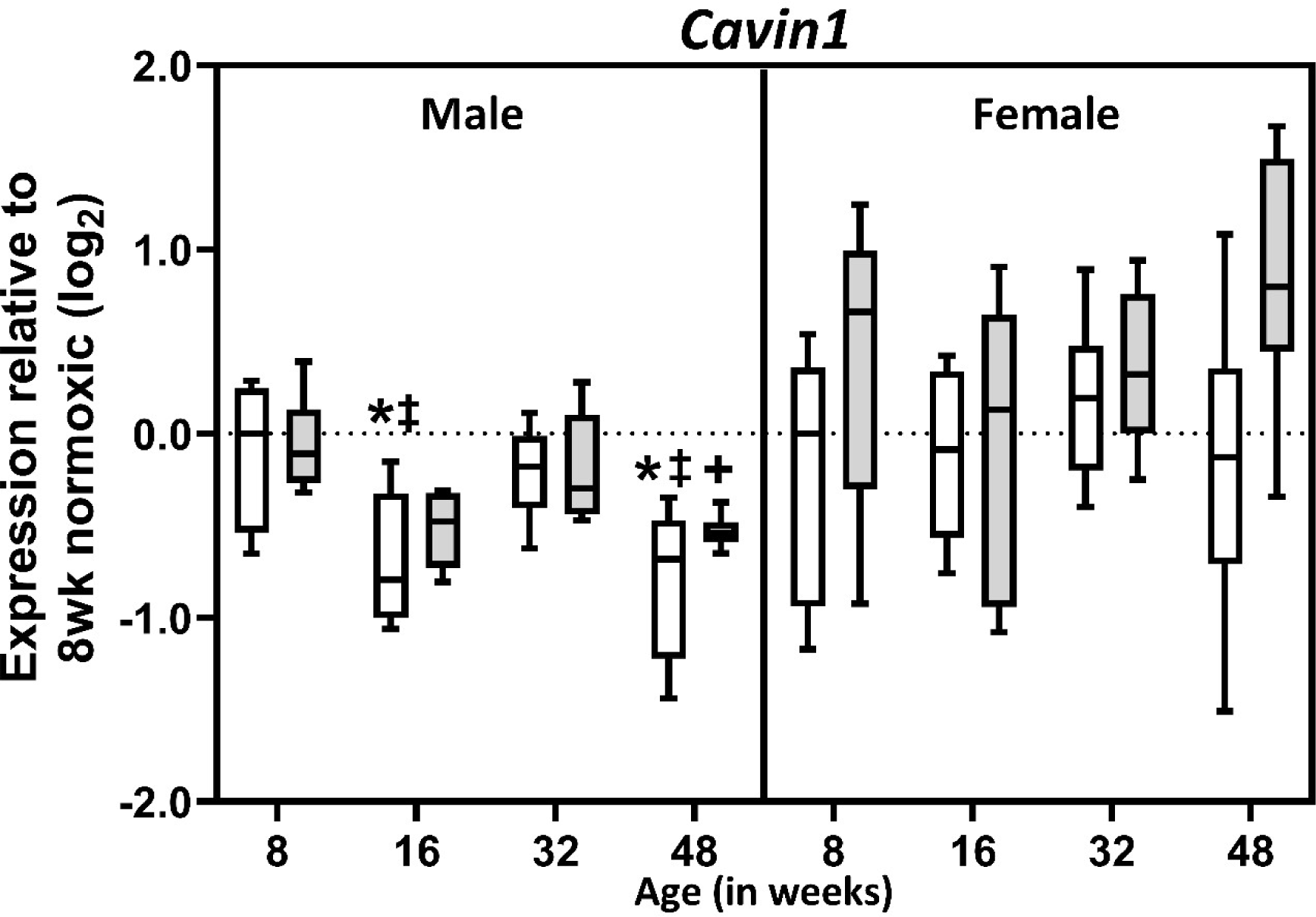
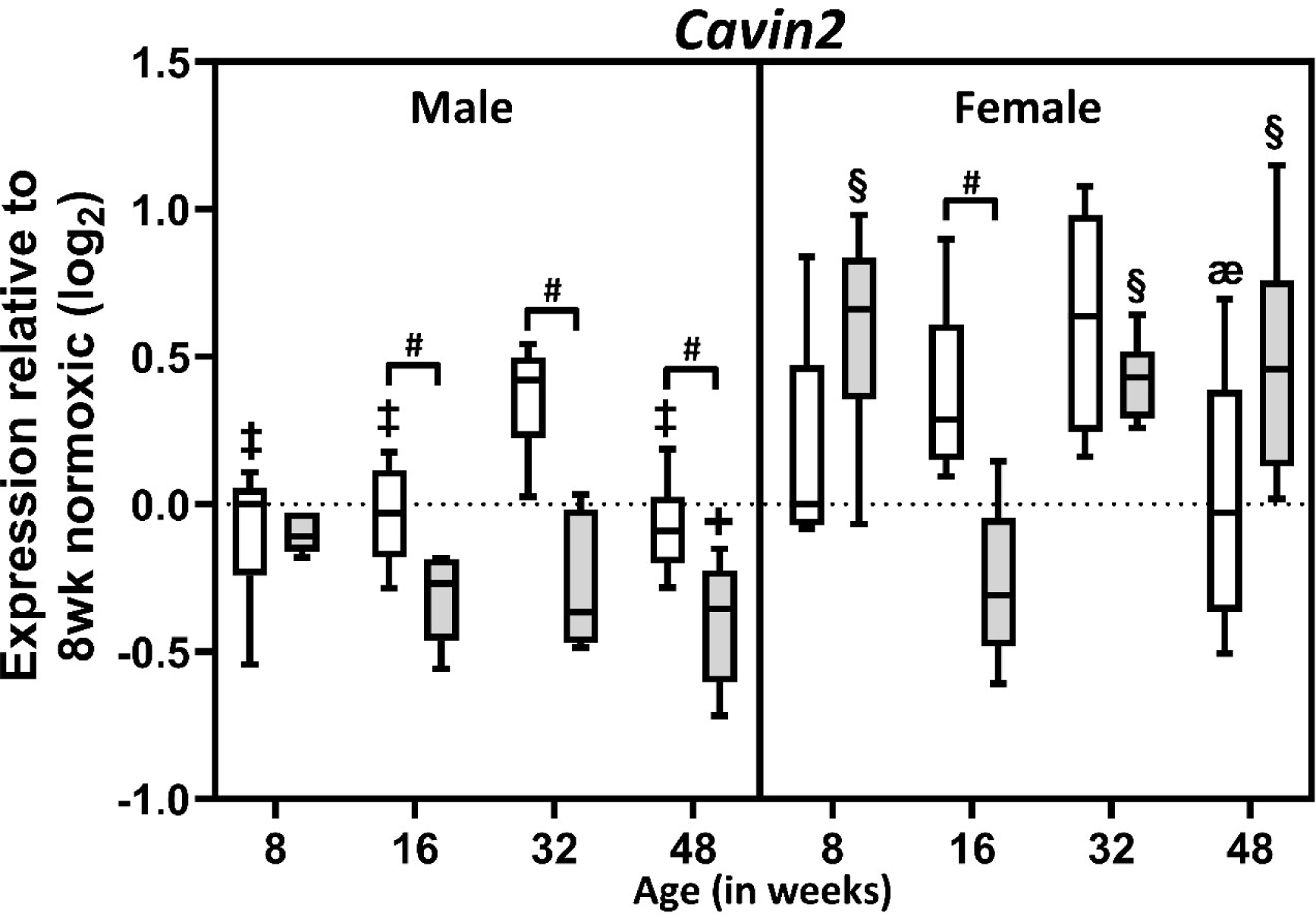
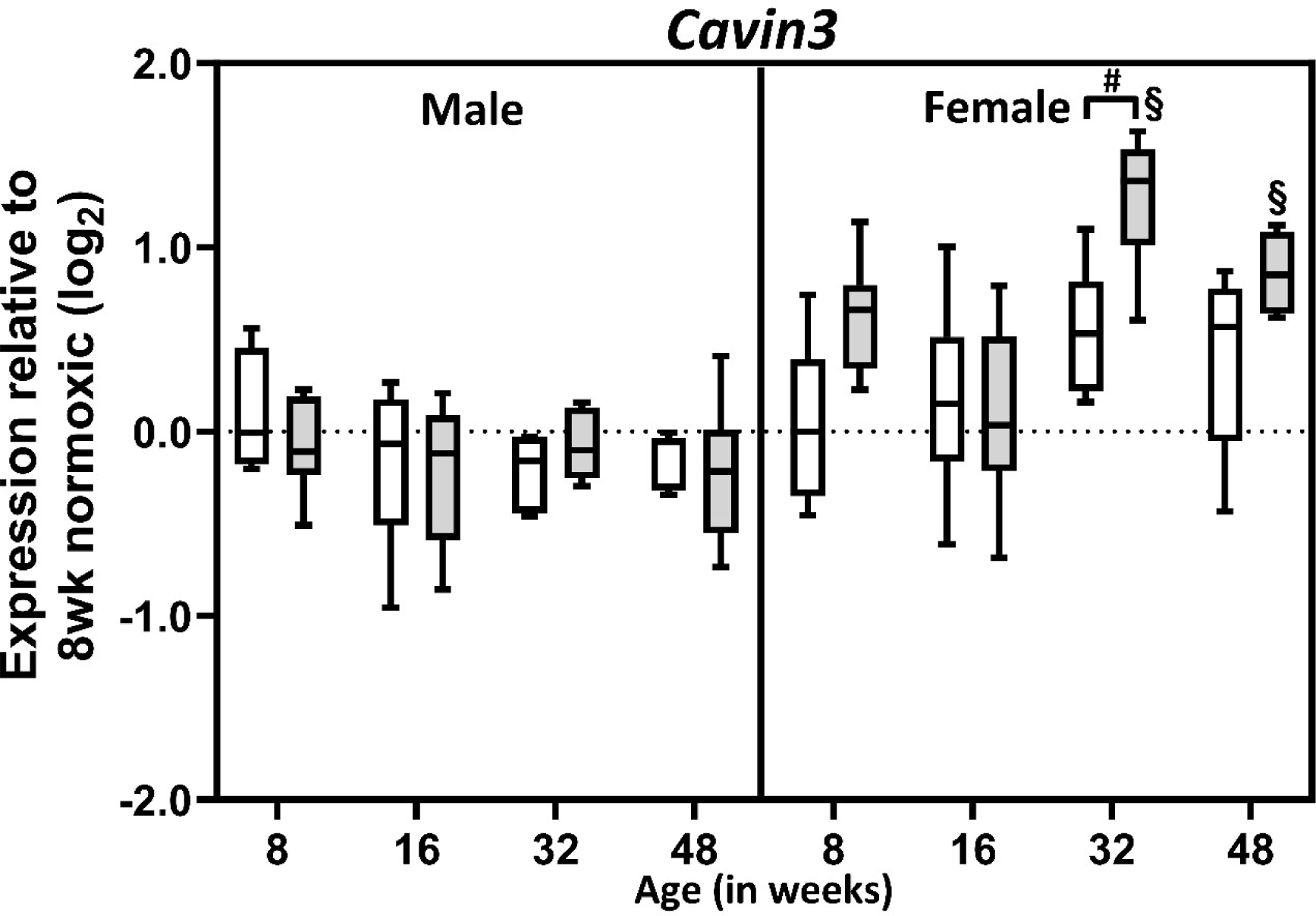
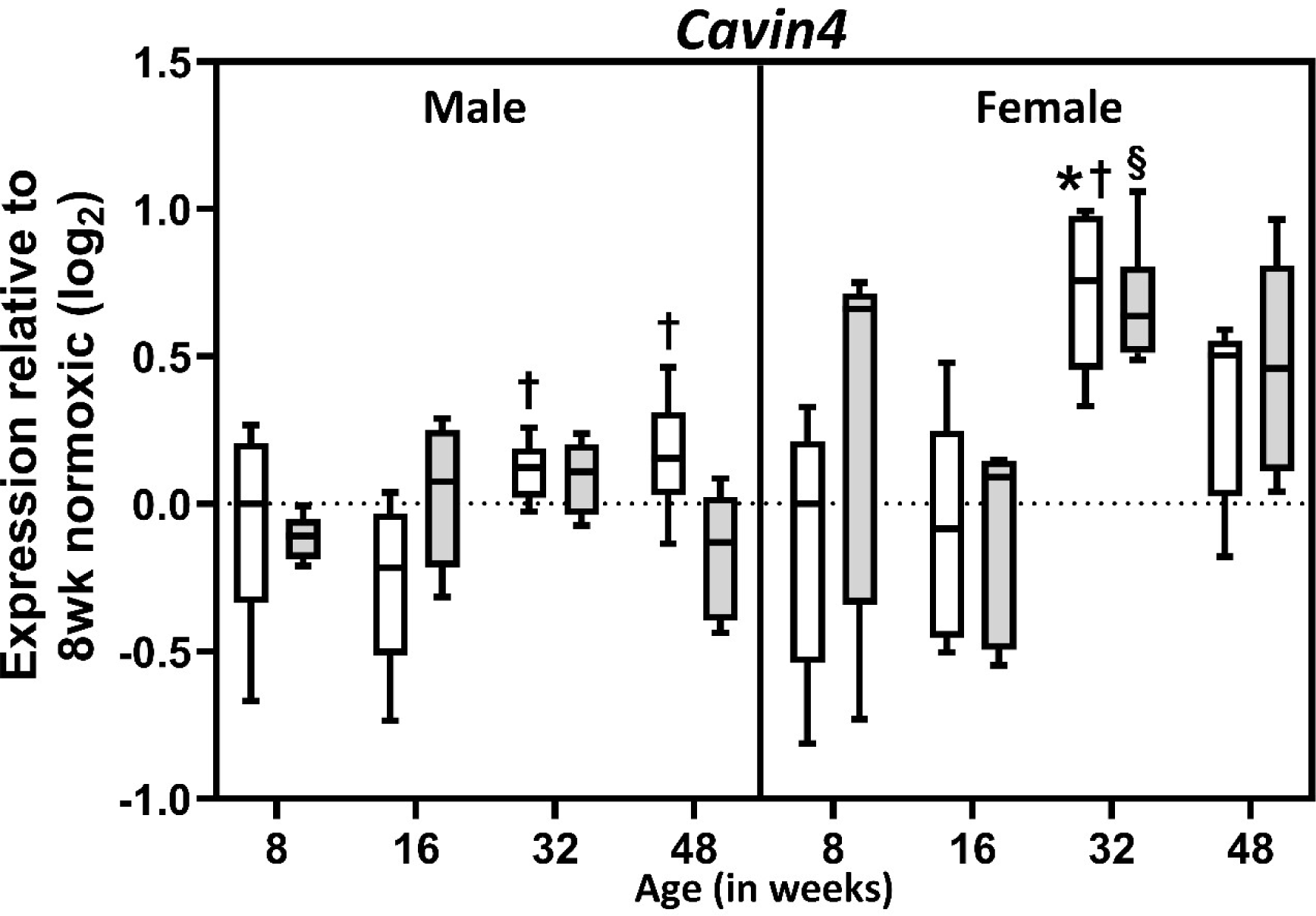
Age-dependent expression of cavin transcripts in normoxic and post-ischemic myocardium. Shown are boxplots detailing expression changes for *Cavin1, Cavin2*, *Cavin3* and C*avin4* transcripts in normoxic (white) and post-ischemic (grey) male and female hearts (*n*=5-6/group). Boxes indicate the interquartile range (25%-75%) with the horizontal bar within each box indicating the median. Whiskers show minimum and maximum values. For normoxic hearts: *, *P*<0.05 vs. 8-week; †, P<0.05 vs. 16-week; ‡, P<0.05 vs. 32 week. For post-ischemic hearts: +, P<0.05 vs. 8-week; §, P<0.05 vs. 16-week; æ, P<0.05 vs. 32-week; #, P<0.05 vs. age-matched post-ischemic group (indicated by brackets).

#### Popdc transcripts

In normoxic hearts, levels of *Popdc1*, *Popdc2* and *Popdc3* transcripts did not vary with age in either males or females (Figure 4). However in post-ischemic male myocardium, aging was associated with a significant 1.4-fold reduction in *Popdc1* (interaction with ischemia; *F*(1,37) = 15.56, *P*=0.0003), and a non-significant reduction (via post-hoc analysis) of *Popdc3* (Figure 4) (interaction with ischemia; *F*(1,35) = 26.20, *P*=0.0001). This trend for *Popdc1* and *Popdc3* was not evident in post-ischemic female hearts. Expression of *Popdc2* was unaffected by age or ischemia in male or female hearts (Figure 4). Detailed two-way ANOVA outputs of caveolin, cavin & Popdc gene expression with cardiac aging are available in the supplementary file table 2 (male) & 3 (female).

**Figure 4.**
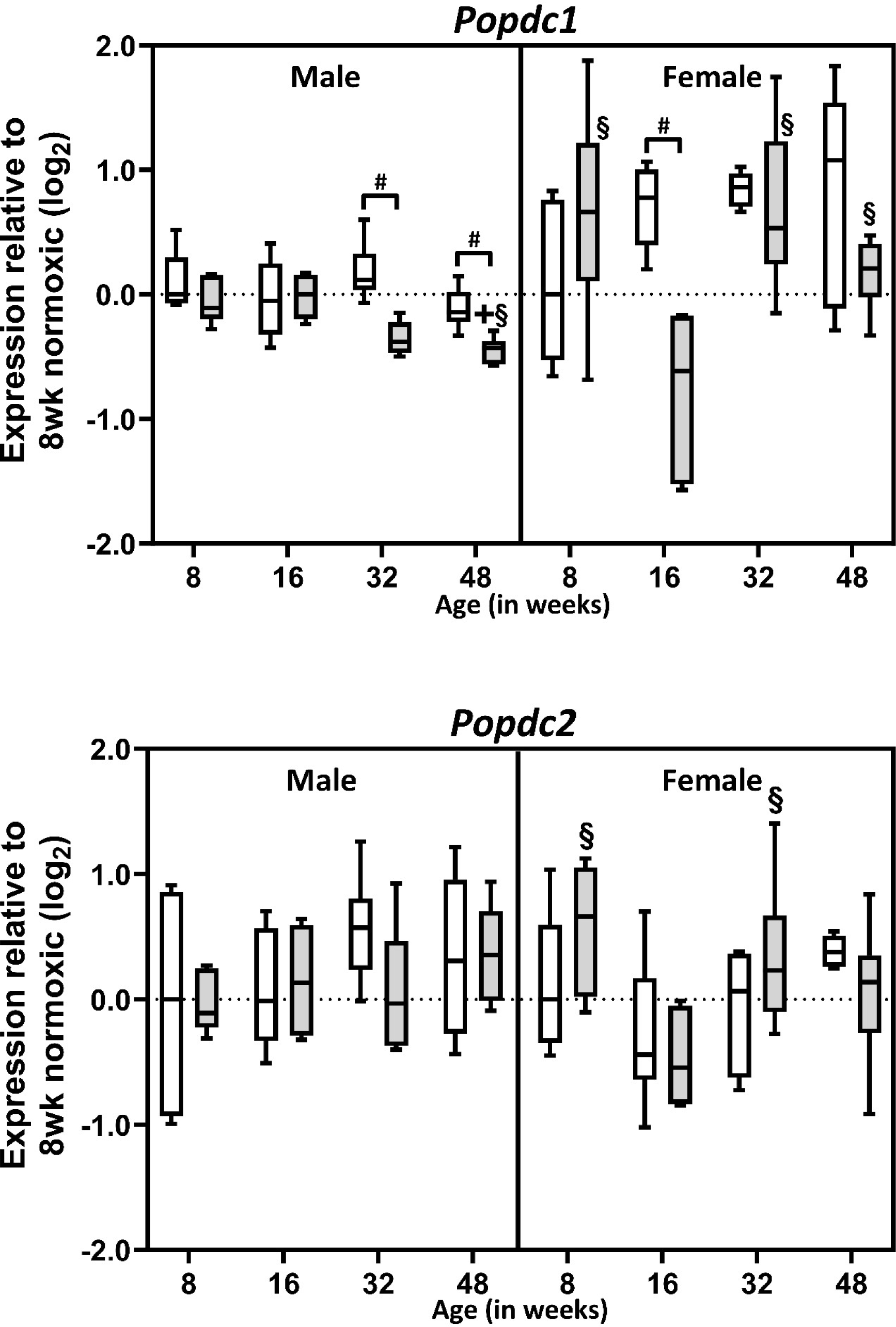
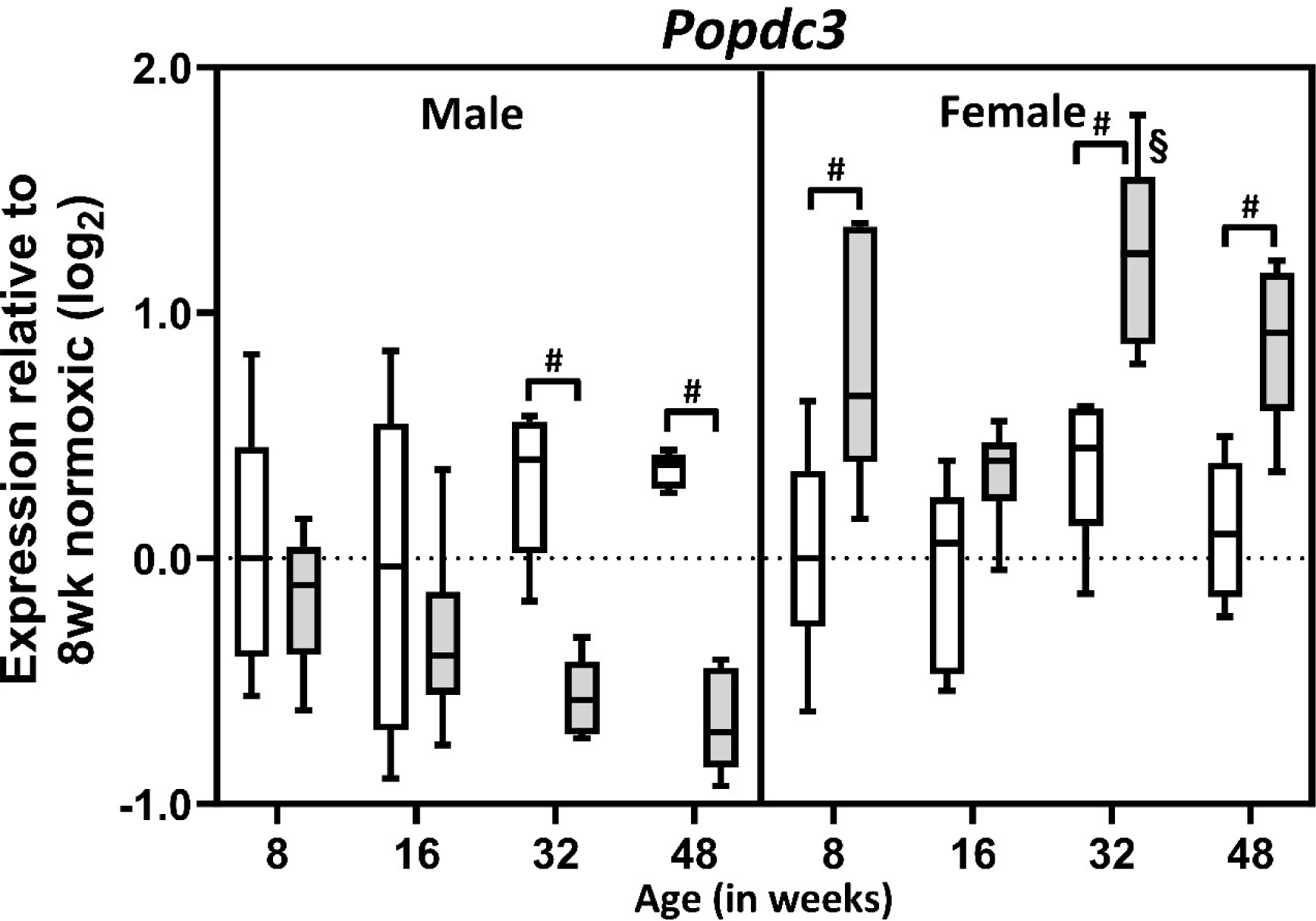
Age-dependent expression of Popeye domain containing transcripts in normoxic and post-ischemic myocardium. Shown are boxplots detailing expression changes for *Popdc1, Popdc2*, and *Popdc3* transcripts in normoxic (white) and post-ischemic (grey) male and female hearts (*n*=5-6/group). Boxes indicate the interquartile range (25%-75%) with the horizontal bar within each box indicating the median. Whiskers show minimum and maximum values. For post-ischemic hearts: +, *P*<0.05 vs. 8-week; §, P<0.05 vs. 16-week; #, P<0.05 vs. age-matched post-ischemic group (indicated by brackets).

#### Predicted & literature screening of microRNA regulators for Cav3 transcript

Using bioinformatics in-silico analysis (Figure 5) supplemented with literature analysis, potential causal microRNA regulators of *Cav3* were screened which may explain age-related changes in *Cav3* gene expression observed in this study. *MiR-22* & -*101b* showed similar expression patterns in both normoxic and post-ischemic male hearts with aging (Figure 6). Despite age-related changes being observed in 48 vs. 8 week old normoxic hearts as well as induction of *miR-22* & -*101b* in 48-week post-ischemic vs. normoxic counterparts, expression patterns of *miR-22* and *miR-101b* did not correlate inversely with *Cav3* transcript expression observed with cardiac aging used in this study as the expression of these microRNAs would be expected to be up-regulated with cardiac aging.

**Figure 5.**
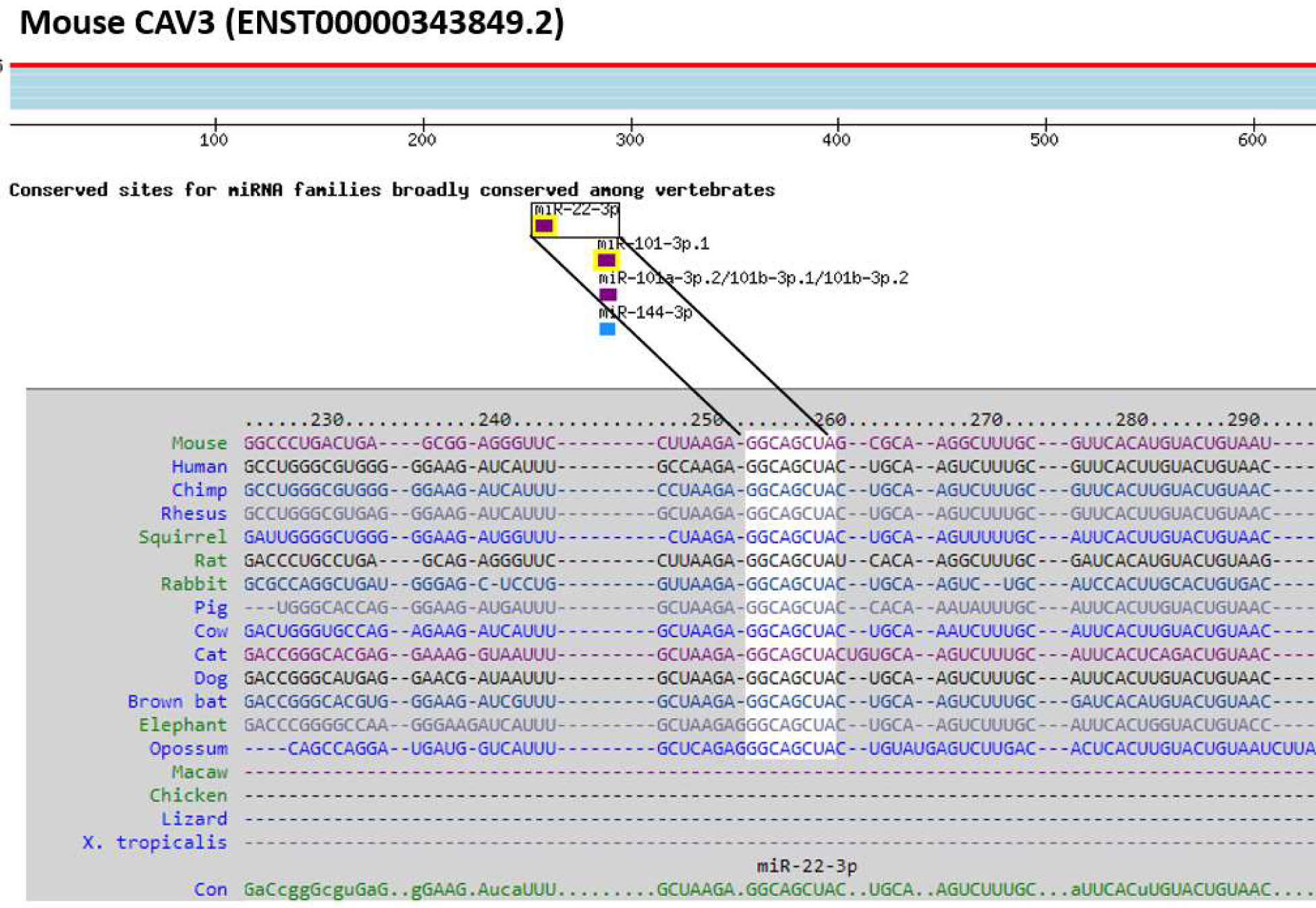
Bioinformatic prediction of mouse microRNA candidates targeting *Cav3* gene using TargetScan. (https://www.targetscan.org/, last accessed: 19-11-2023).

**Figure 6.**
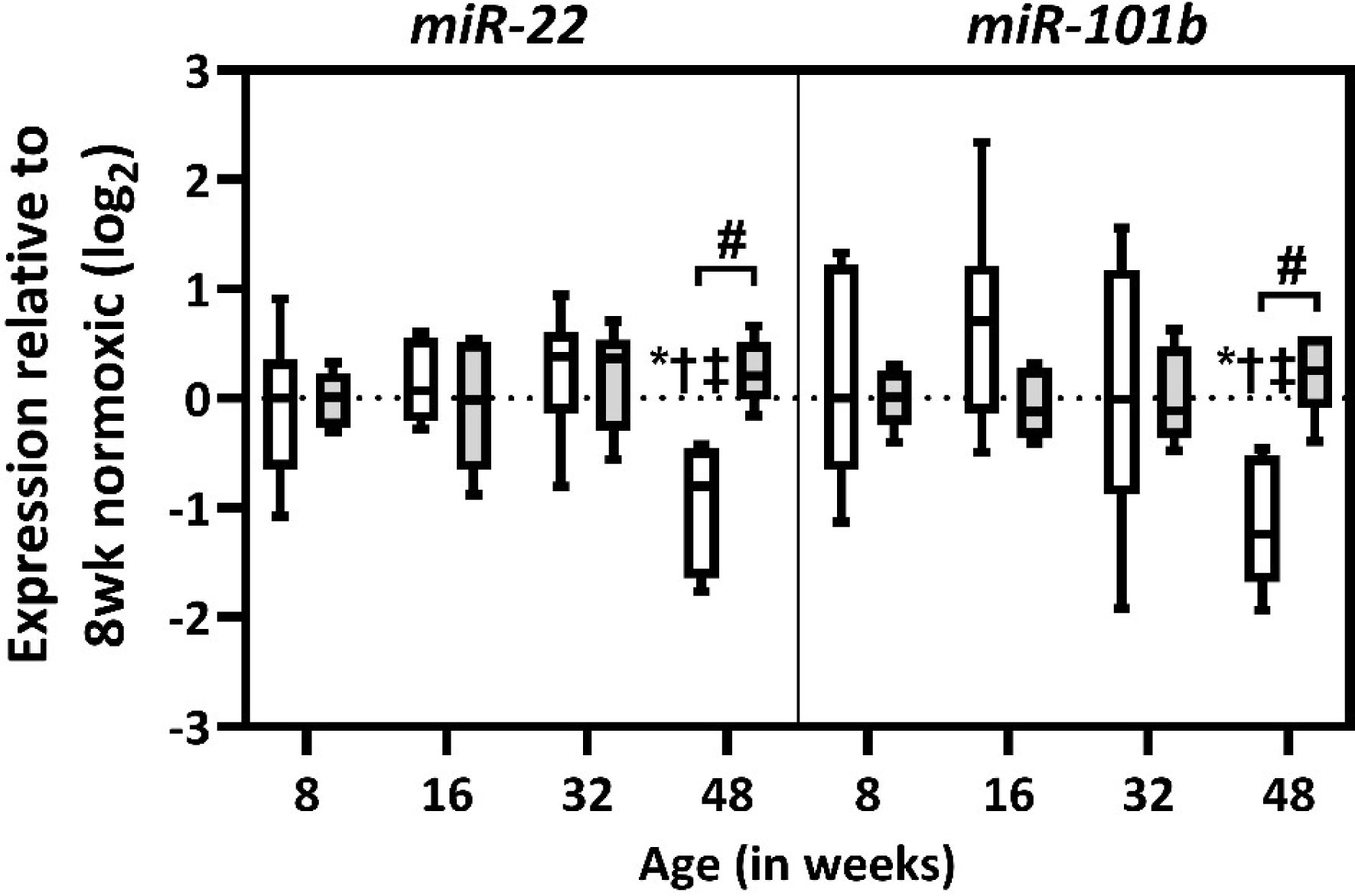
Age-dependent expression of microRNA-22 & microRNA-101b transcripts in normoxic and post-ischemic male myocardium. Shown are boxplots detailing expression changes for *miR-22* (left) & *miR-101b* (right) transcripts in normoxic (white) and post-ischemic (grey) male hearts (n=5-6/group). Boxes indicate the interquartile range (25%-75%) with the horizontal bar within each box indicating the median. Whiskers show minimum and maximum values. For normoxic hearts: *, P <0.05 vs. 8-week; †, P<0.05 vs. 16-week; ‡, P<0.05 vs. 32-week. For post-ischemic hearts: #, P<0.05 vs. age-matched post-ischemic group (indicated by brackets).

#### Cav-3 and Cavin-1 protein expression in the aging heart

In agreement with transcriptional analysis, analysis of Cav-3 protein in 48 vs. 8-week male hearts revealed modest reduction (by 38%) in normoxic expression (Figure 7) while Cavin-1 showed 52% reduction. Also congruent with transcriptional analysis, Cavin-1 protein levels did not exhibit age-dependence in normoxic & post-ischemic female myocardium (Figure 8).

**Figure 7.**
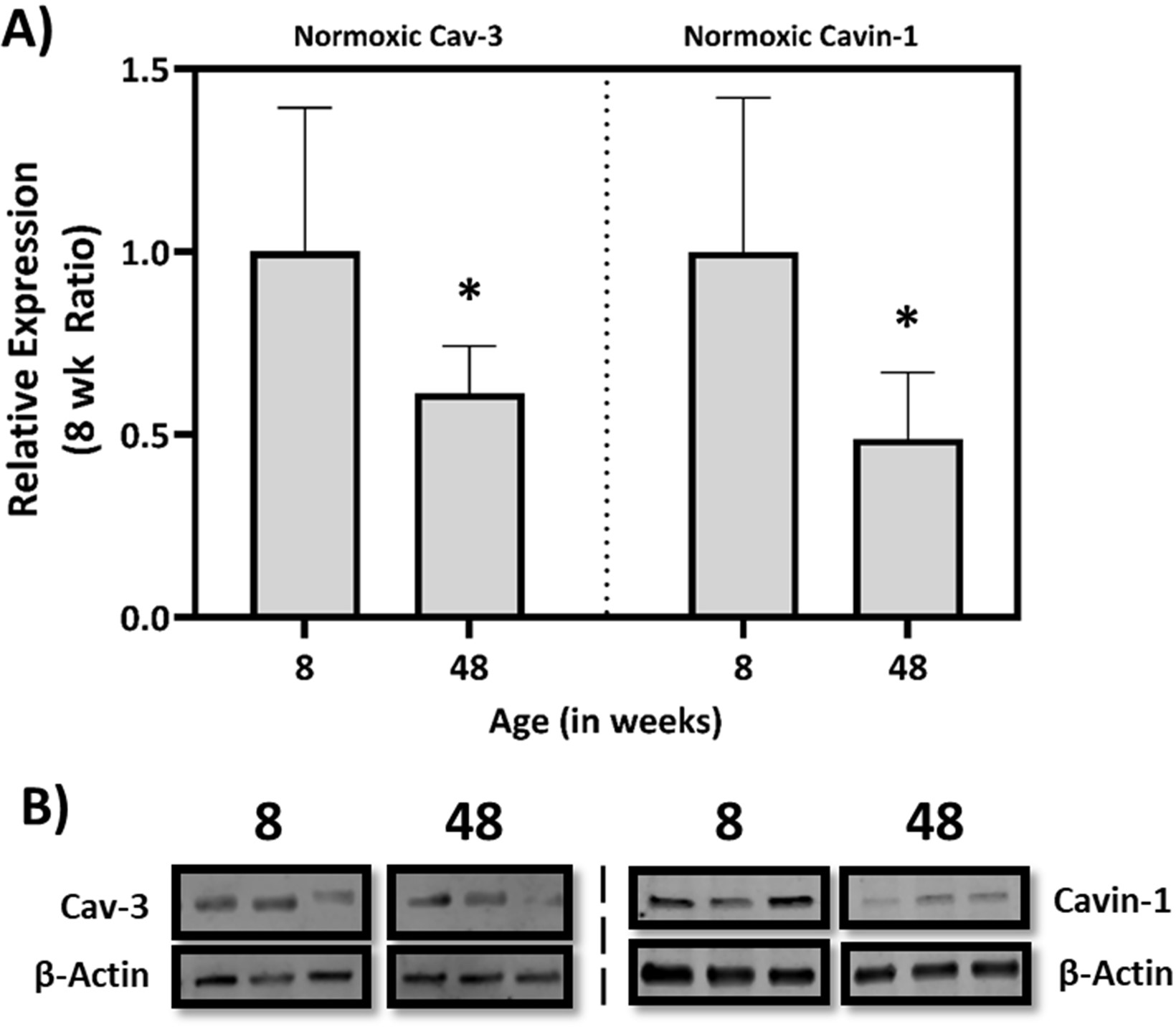
Age dependent expression of Caveolin-3 & Cavin-1 protein in the normoxic male myocardium. **A)** Bar graphs present expression levels for left ventricular Caveolin-3 & Cavin-1 protein in normoxic male hearts (*n*=6/group). **B)** Immunoblot confirmation of normoxic Cav-3 (left inset) & Cavin-1 (right inset). Data are expressed as mean ± SEM. *, *P*<0.05 vs. 8-week.

**Figure 8.**
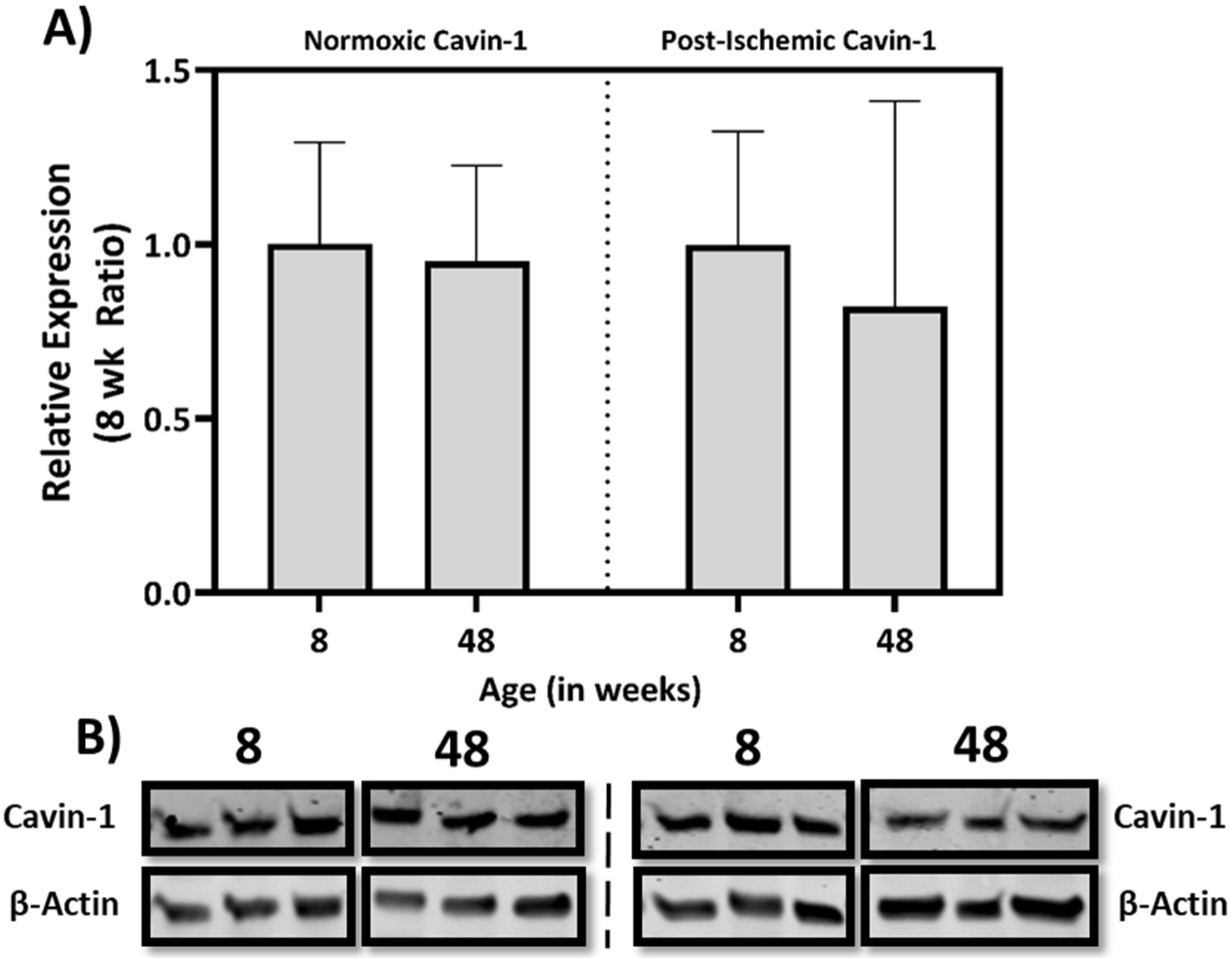
Age dependent expression of Cavin-1 protein in the female normoxic and post-ischemic myocardium. **A)** Bar graphs detailing expression changes for left ventricular Cavin-1 protein in normoxic and post-ischemic female hearts (*n*=6/group). **B)** Immunoblot confirmation of normoxic Cavin-1 (left inset) & post-ischemic Cavin-1 (right inset). Data are expressed as mean ± SEM. *, *P*<0.05 vs. 8-week.

### Simulated ischemia-reperfusion in HL-1 cardiomyocytes

As shown by LDH efflux, 3 hour ischemia followed by 5 hours of reperfusion significantly induced cellular injury relative to normoxic cells in HL-1 cardiomyocytes (Figure 9). Pretreatment with 100 nM of selective A_1_AR agonist CCPA resulted in a cytoprotective effect by lowering LDH release (by approx. 20%) when compared to simulated I-R group (Figure 9). This cardioprotective effect of A_1_AR agonism was abolished when CCPA incubation was combined with 1 mM of membrane cholesterol depleting agent MβCD which also reduced ischemic tolerance evidenced by enhanced LDH release in this group when compared to simulated I-R group (Figure 9). More specifically, pretreatment of MβCD with CCPA resulted in 20% more LDH release when compared to simulated I-R group.

**Figure 9.**
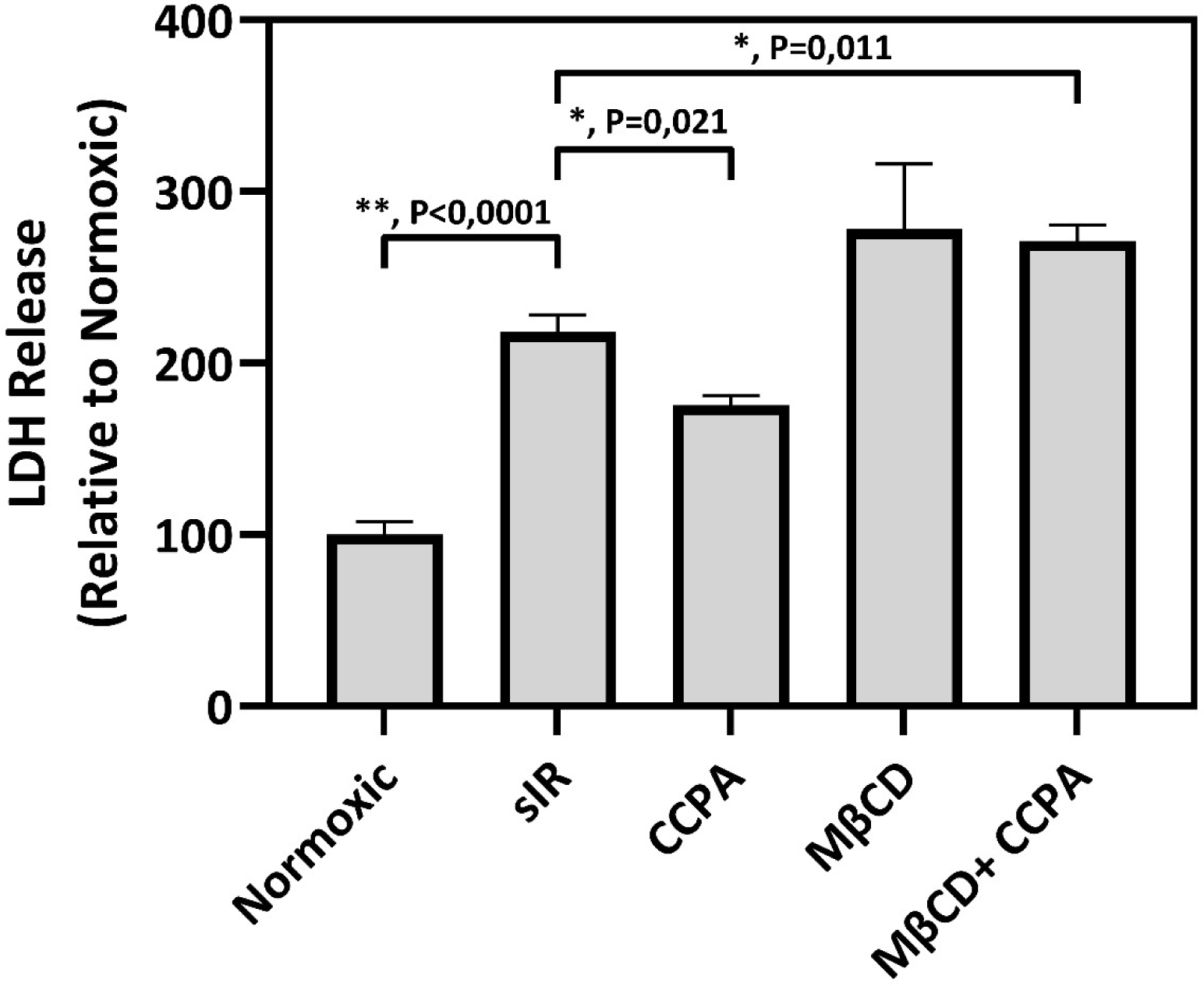
LDH release following acute A_1_AR agonism (via CCPA) and caveolar distruption (via MβCD) in HL-1 cardiomyocytes subjected to simulated ischemia-reperfusion. Data are expressed as mean ± SEM (n=6). *, *P*<0.05 vs. sIR, **, P<0.0001 vs. sIR.

## Discussion

Aging is associated with reductions in both myocardial ischemic tolerance and the efficacy of cardioprotective stimuli, including ischemic preconditioning and GPCR-mediated protective responses (Peart et al., 2014; Schilling and Patel, 2016). Caveolae and their coat proteins are critical to responses to both injurious and protective stimuli, localizing or modulating cardioprotective receptors as well as G proteins, its downstream effector molecules (Stary et al., 2012; Head et al., 2005; Schilling et al., 2018), autophagy (Kassan et al., 2016), as well as being critical to contractility by maintaining t tubular I_Ca_ (Bryant et al., 2018; Kong et al., 2019). Numerous studies confirm that caveolar disruption via chemical methods or genetic ablation of *Cav3* negates the effects of multiple cardioprotective stimuli and attenuates intrinsic ischemic tolerance (Tsutsumi et al., 2008; See Hoe et al., 2014; Horikawa et al., 2008; Tsutsumi et al., 2010a; Tsutsumi et al., 2010b), conversely cardioprotective interventions such as δ-opioid preconditioning increases caveolae formation and caveolin expression in the heart (Wang, Xue and Wu, 2020). Similarly, previously was shown in *in vitro* cultured cardiomyocytes A_1_AR agonism via acute CCPA administration is cytoprotective and the conditioning response is abolished by MβCD treatment, similar to previous *ex vivo* observations for acute opioid receptor conditioning (See Hoe et al., 2014). Thus the age- and sex-dependence of transcription and expression of major caveolar elements (caveolin, cavin and Popdc families) was characterized in the normoxic and post-ischemic myocardium.

Results highlight age-related impairment of I-R tolerance in male and female hearts, similar to a previous study which assessed ischemic tolerance in the aging heart (Willems et al., 2005). In the male hearts, stress-intolerant middle-aged hearts were associated with disruption of caveolae-related transcripts (and proteins), specifically with shifts in *Cav1, Cav3 Cavin1, Cavin2*, *Popdc1* and *Popdc3* transcripts in male hearts. These data are consistent with reports of age-related reductions (∼38%) in myocardial Cav-3 and caveolae in 74-week-old male “aged” hearts (Peart et al., 2014; Kong et al., 2018), old pacemaker cells (Choi et al., 2022), together with age-associated reductions in caveolar proteins in other tissues, including bladder smooth muscle (Lowalekar et al., 2012), urothelium of the bladder (Kim, Yu and Kim, 2021), skeletal muscle (Barrientos et al., 2015), and neurons (Head et al., 2010). There is also evidence for sarcolemmal disappearance of Cav-3 and reductions in cardiac caveolins, cavins and Popdc in human and rat heart failure (Ratajczak et al., 2003; Feiner et al., 2011; Gingold-Belfer et al., 2011; Norman et al., 2014; Colak et al., 2016). In contrast to males, chief caveolar coat protein genes *Cav3*, *Cavin1* and *Popdc1* were largely stable with cardiac aging in females. To best of our knowledge, this was the first study that attempted to elucidate the expression profiles of chief caveolar coat genes in the aging female heart.

### Caveolins

Of the caveolin transcripts only *Cav3* was consistently down-regulated with age in male hearts, translating to 38% reductions in normoxic Cav-3 expression. These data support the notion that reductions in Cav-3 (and caveolae) may be important mechanistic determinants of ischemic intolerance and refractoriness to protective stimuli with aging (Peart et al., 2014). Experimental depletion of caveolae & Cav-3 significantly reduces endogenous I-R tolerance, pharmacological and ischemic preconditioning responses, while its increased expression promotes stress tolerance (Kassan et al., 2016; Fridolfsson et al., 2012; Tsutsumi et al., 2008; Garcia-Nino et al., 2017). Age-related repression of *Cav3* mRNA and protein, together with shifts in other caveola-related transcripts, is consistent with reports of reduced myocardial Cav-3 protein and caveolae density in aged (74-week) mouse hearts observed by other authors (Peart et al., 2014; Kong et al., 2018). Age may also suppress baseline (normoxic) expression of *Cav1* transcript, with Cav-1 also implicated in promoting myocardial ischemic tolerance as well as cardiac cell elasticity, contractility and pumping efficiency which could explain the significant reduction in cardiac performance (Jasmin et al., 2011; Grivas et al., 2020). However, effects of its down-regulation appear to differ from Cav-3 depletion, with a select reduction in contractile recovery without alterations in infarct size, in association with reductions in post-ischemic β-adrenoceptor, cAMP and phospho-AKT levels (Jasmin et al., 2011). Besides this, a recent study has also shown a short peptide moiety delivery of Cav-1 containing Caveolin Scaffolding Domain reduces aging-associated increase in cardiomyocyte cross-sectional area and enhanced ventricular compliance evidenced by echocardiographic studies, leading to improved ejection fraction, fractional shortening and reduced isovolumic relation time (Kuppuswamy et al., 2021).

In mouse intestinal I-R, Cav-1 has been shown to restrain excessive PKCβII activation, further inhibiting oxidative stress and apoptosis via the PKCβII/p66shc pathway (Chen et al., 2023) with Cav1-deficiency following cerebral I-R also shown to result in increased volume of infarction (Jasmin et al., 2007). Additionally, Cav-1 peptide delivery using the Caveolin Scaffolding Domain for 4 weeks has been shown to reverse pathologic changes such as microvascular leakage & fibrosis in all organs similar to the levels in young healthy mice (Kuppuswamy et al., 2021). We also noticed a decline in expression of *Cav2* with aging in the normoxic male heart (but not females), previously Cav-2 overexpression has been shown to major be a regulator of longevity and stress adaptation in *C. elegans* (Mehta et al., 2016). In an aging model of the mouse brain, *Cav2* transcript & protein was found to be up-regulated which was associated with age-related neuroinflammation (Park et al., 2022). In contrast, a recent study suggested a protective role of *Cav2* in an intestinal model of I-R injury where infiltrating cytotoxically activated leukocytes were found in *Cav2*-deficient mice subjected to I-R which were associated with enhanced leukocyte rolling, adhesion and injury (Liu et al., 2021). Collectively, suppression of Cav-1, -2 and -3 is predicted to significantly impair the hearts ability to withstand or adapt to ischemic insult with age.

### Cavins and Popdcs

We also observed significant age-dependent reductions in Cavin-1 transcript and protein expression in male hearts, while Cavin-1 protein levels were stable in female normoxic & post-ischemic hearts. Although Cav-3 has been more widely studied for its role in stress responses and cardioprotection, Cavin-1 and Popdc1 are also important to cardiomyocyte caveolae formation and I-R tolerance (Alcalay et al., 2013; Taniguchi et al., 2016; Kaakinen et al., 2017). Cavin-1 is also implicated in plasma membrane repair, via interaction with plasma-membrane repair machinery such as Trim72/Mg53 and Smpd1, with *Cavin1* deletion reducing membrane resealing and promoting cell death (Corrotte et al., 2013; Zhu et al., 2011), consistent with the findings that caveolins and cavins play a crucial role in sarcolemmal mechanoprotection (Lo et al., 2015). Cavin-1 also stabilizes caveolins as *Cavin1*-deficiency has been shown to reduce cardiac caveolin expression and cardiomyocyte caveolae (Taniguchi et al., 2016; Liu et al., 2008; Kaakinen et al., 2017). Cavin-1 has also been shown to be a crucial determinant of sarcolemmal fragility and myocardial responses to stretch and ischemia (Kaakinen et al., 2017), and Cavin-1 and -3 have been shown to influence cell survival/death signaling (McMahon et al., 2019) as well as autophagy (Kassan et al., 2016).

We observed interaction of age and ischemia for *Cavin2* in the aging male myocardium more specifically repression of *Cavin2* in post-ischemic middle-aged hearts. Though yet to be shown in the heart (specifically in cardiomyocytes), Cavin-2 reduction via knockdown has been shown to result in detrimental outcomes in lung models of I-R injury, conversely *Cavin2*-overexpresison has been found to reduce production of inflammatory factors and oxidative stress injury (Kasahara et al., 2023; Tang et al., 2022). Similarly a recent report suggests that fibroblast-specific Cavin-2-deficiency attenuates cardiac fibrosis resulting in the suppression of heart failure (Higuchi et al. 2024). We also observed up-regulation of *Cavin3* in the aging I-R female hearts, its role on myocardial caveolae formation and I-R signaling remains unknown at the time of this writing. Cavin-4 through its coiled-coil domain has been suggested to stabilize Cav-3 at the plasma membrane of cardiomyocytes (Naito et al., 2015). It is unclear whether Cavin-4 deficiency would result in caveolae reduction; its deletion has been shown to reduce infarct size after cardiac I-R and preserved cardiac contraction (Nishi et al., 2019). In the present study, expression patterns of *Cavin4* transcript were complex with aging, ischemia and male vs. female.

It is unknown whether Popdc1-deficiency impacts caveolin and cavin expression: Popdc1-deficient hearts display a shift from Cav-3 rich fractions towards a lower buoyant density, suggesting alterations in caveolar makeup in these hearts (Alcalay et al., 2013). Popdc1-deficient hearts also display refractoriness to protective effects of ischemic and anesthetic preconditioning, similar to Cav-3-deficiency (Alcalay et al., 2013; Horikawa et al., 2008; Tsutsumi et al., 2010a), while small interference RNA-mediated gene silencing shown to result in cardiomyocyte injury and death (Kliminski, Uziel and Kessler-Icekson, 2017). Disruption of Popdc1 and Popdc2 results in stress-induced sinus node dysfunction, chronotropic incompetence and long sinus pauses, with evidence of age-dependent maladaptive cardiac pacemaking responses to stress (swimming/exercise) (Froese et al., 2012). Furthermore, variants in the *POPDC1* gene such as *POPDC1^S201F^* show high creatine kinase levels in their blood which may be due to impaired membrane repair response (Brand, 2019). While normoxic or baseline *Popdc* transcript levels were not suppressed with age in males or females (indeed, *Popdc1* and *Popdc3* increased in older females and males, respectively), age-dependent reductions in post-ischemic *Popdc1* and *Popdc3* transcripts were evident in male hearts. It remains to be determined whether these shifts in Popdc transcripts translate to shifts in protein expression.

### Sex-related effects

Expression of caveolar transcripts/proteins in males and females were characterized as epidemiologic and experimental evidence indicate female hearts have greater resistance to I-R injury and ischemic heart disease development compared with age-matched males (Willems et al. 2005; Ostadal and Ostadal, 2014; Yusifov, Woulfe and Bruns, 2022). Menopause is thought to reverse this cardioprotection, consistent with involvement of sex hormones. Sex-based studies have largely focused on estrogen receptors α, β and G protein-coupled receptor 30, expressed in both male and female myocardium. While activation of these receptors is cardioprotective there is conflicting evidence as to whether they are caveolae-dependent and whether Cav-3 and estrogen receptor α co-localize in cardiomyocytes (Chambliss et al., 2000; Chung et al., 2009; Mahmoodzadeh et al., 2006) though there is evidence of co-localization in other cell types (Wong et al., 2019). Estrogen stimulation (i.e. by E2) also stimulates Cav-1 and Cav-2 protein synthesis which are also required for proper targeting of ER-α to the membrane (Razandi et al., 2002; Evinger and Levin, 2005; Totta et al., 2016). Ovariectomy results in a decrease in Cav-3, and estrogen receptor agonism (i.e. by E2) up-regulates Cav-3 protein (Cui et al., 2011; Tan et al., 2012). Sex-related differences in eNOS and nNOS translocation in stressed myocardium (with I-R or Ca^2+^ loading) have also been observed, with females exhibiting enhanced eNOS and nNOS association with Cav-3 (Sun et al., 2006).

There are conflicting reports on sex-related differences in ischemic tolerance with aging: some investigators have reported no sex differences in ischemic tolerance (Sun et al., 2006; Cavasin et al., 2004). Interestingly, we do observe a broader impact of age on transcription and expression of caveolae proteins in males vs. females. Normoxic female hearts exhibited more stable patterns of caveolin, cavin and Popdc transcript expression, whereas crucial caveolae forming proteins such as Cav-3 and Cavin-1 were down-regulated with age in the normoxic male hearts. Deletion of Cavin-1 has been has been shown to result in loss of myocardial caveolae and exacerbation of post-ischemic contractile dysfunction and cell damage (Kaakinen et al., 2017). In addition, Cavin-1 deficiency has been observed to result in a sex-dependent fall in cardiac function *in vivo* in females (∼20% lower ejection fraction), suggesting greater dependence on Cavin-1 in female hearts (Reichelt et al., 2016). Though not assessed here, a similar aging study suggest sex-related differences in tolerance to I-R injury for middle-age female vs. male hearts (Willems et al., 2005) with females shown to have increased ischemic tolerance in young, adult and middle-aged hearts vs. male counterparts. As sarcolemmal caveolae-forming genes such as *Cav3*, *Cavin1* and *Popdc1* are largely stable in the aging female normoxic & post-ischemic heart, our results align with aforementioned findings of enhanced ischemic tolerance (evidenced by LDH efflux) in female hearts as caveolae are crucial determinants of stress tolerance (See Hoe et al., 2014; Horikawa et al., 2008; Peart et al., 2014).

### Transcriptional regulation via miR-22 & 101b

As *Cav3* mRNA was consistantly down-regulated with aging in the male myocardium (to a lesser degree in females), we hypothesized whether this transcriptional repression could be related to microRNA regulation of gene expression as seen by others with cardiac aging and disease (Boon et al., 2013; Gatsiou et al., 2021; Bei et al., 2018). In-silico bioinformatic analysis of *Cav3* and literature search (Chen et al., 2015; Li et al., 2021; Zhang et al., 2018) for *Cav3*-related microRNAs revealed *miR-22* & *miR-101b* as top candidates for regulation of *Cav3*. Although *miR-22* and *miR-101b* showed significant age-dependant changes in the normoxic heart, their expression pattern did not (inversely) correlate with *Cav3* transcriptional changes seen in the normoxic and post-ischemic male hearts. It is possible that *Cav3* transcript expression is controlled by other microRNAs which were not assessed here, namely those of miR-101/101a family, miR-144 and other types of gene regulators (i.e. transcription factors, long non-coding, circular RNAs) or microRNAs yet to be discovered (Ghafouri-Fard et al., 2022). Indeed, hypoxia-inducible factor-1α has been shown to be a causal mechanisms for down-regulation of caveolin, cavin expression & caveolae formation in response to hypoxia in 3T3-L1 adipocytes & human (hMADS) adipocytes (Regazzetti et al., 2015; Varela-Guruceaga et al., 2018). However in a similar previous study to the one here with cardiac aging hypoxia-inducible factor-1α was not differentially expressed (Ashton et al., 2023).

### Conclusions

The current study evidences reductions in caveolae-related transcripts and proteins in middle-aged hearts, correlating with declining ischemic tolerance in males but not females. In the male myocardium, suppression of Cav-3 and Cavin-1 expression, together with shifts in transcription of *Cav1*, *Cav3*, *Cavin1*, *Cavin2*, *Popdc1* and *Popdc3* may impair caveolae formation/function and caveolae-dependent stress signaling (See Hoe et al., 2014; Kiessling and Ashton, 2017). These transcriptional changes are more prominent in male vs. female hearts, which may be relevant to sex-dependent differences in stress responses, though this requires further study. Indeed, whether changes in other cavin and Popdc transcripts consistently translate to altered protein expression awaits further study (as does the impact of altered Cavin-2 and Popdc3 expression on stress tolerance). Besides the contribution of transcriptional shifts resulting in altered protein expression, we hypothesize whether observed proteomic changes in expression has subsequent effects on other caveolar coat proteins. For example *Cav1*-deficiency in the heart has been shown to significantly reduce Cav-2, Cav-3, and Cavin1-4 (Hansen et al., 2013). Similarly, *Cavin1*-deficiency has been shown to negatively affect caveolin and other members of cavins with proteosomal degredation shown to be a mechanism of Cavin-1 protein inhibition (Hansen et al., 2013; Kaakinen et al., 2017; Taniguchi et al., 2016; Zhou et al., 2017).

### Future Directions

Future studies should additionally characterize expression of other protein families influencing caveolae formation/function, such as dynamins, EHD, pacsin and possible cytoskeletal effectors which help coordinate mechanoadaptation via stress fiber remodeling (Han et al., 2016; Echarri et al., 2019). As many protective stimuli or interventions are reliant on caveolae, unraveling mechanisms governing the age-dependencies of caveolin, cavin and Popdc transcription and expression may facilitate targeted modulation of this important system. For example, intracoronary adenoviral *Cav3* gene delivery may be a useful experimental approach to enhancing resistance to the damaging impacts of ischemic heart disease, as it has been shown to markedly improve caveolae formation and recovery in the aged heart (Kidd et al., 2010). Similarly, a recent preliminary study has shown that adeno-associated virus delivery of *Cavin1* can enhance the expression of *Cav1*, *Cav3*, *Cavin1* and *Cavin2* transcripts (Russell et al., 2021) which showed age-related changes in this study. What mechanisms govern transcriptional changes with aging also requires further study.

## Acknowledgments

This work contains biobank samples & material contributions supported by grants from the National Health and Medical Research Council of Australia (628841, 481922), Griffith University and Bond University. Would like to thank Dr. Kevin Ashton for assistance in manuscript preparation and experiment design, as well as Dr. Louise See Hoe and Dr. Boris Budiono for their technical support with immunoblotting.

## Acronyms

A_1_AR: Adenosine Receptor Subtype 1
Cav-1 (protein): Caveolin-1 (protein)*, Cav1* (mRNA)
Cav-2 (protein): Caveolin-2 (protein), *Cav2* (mRNA)
Cav-3 (protein): Caveolin-3 (protein), *Cav3* (mRNA)
CCPA: 2-chloro-N6-cyclopentyadenosine
GPCR: G-protein coupled receptor
I-R: Ischemia Reperfusion
LDH: Lactate Dehydrogenase
LVDP: Left Ventricular Developed Pressure
MβCD: Methyl-β-cyclodextrin
miR: miRNA - microRNA
Popdc1 (protein): Popeye domain-containing 1 (protein), *Popdc1* (mRNA)
Popdc2 (protein): Popeye domain-containing 2 (protein), *Popdc2* (mRNA)
Popdc3 (protein): Popeye domain-containing 3 (protein), *Popdc3* (mRNA)
sIR: simulated ischemia-reperfusion

