## Supplementary data for "Age- and Sex-Dependent Caveolar Gene Expression in Normoxic and Post-Ischemic Mouse Hearts"

**Supplementary Table 1. RT-qPCR primer sequences.**

| Protein Name | Gene symbol(s) | NCBI GeneID | Forward primer (5'-3')<br>Reverse primer (5'-3') |
|---|---|---|---|
| Caveolin-1 | *Cav1* | 12389 | AGGTGACTGAGAAGCAAGTG<br>TCAAAGTCAATCTTGACCACG |
| Caveolin-2 | *Cav2* | 12390 | TTGACTACGCAGATCCTGAG<br>GAAGCCTAGCTTGAGATGAG |
| Caveolin-3 | *Cav3* | 12391 | GGATCTGGAAGCTCGGA<br>AATCTACCTTCACAATGTCCTC |
| Cavin1 | *Cavin1/ Ptrf* | 19285 | GATCTACCAGGATGAAGTCAAGC<br>CTCGTCACCCTCCTTCTCAG |
| Cavin2 | *Cavin2/ Sdpr* | 20324 | GCTCATCTTCCAGGAAGAAAGTG<br>AGGGACTTGTTCTCATCAGC |
| Cavin3 | *Prkcdbp/ Cavin3* | 109042 | TGCTCTTCAAGGAGGAGACTG<br>CATCCTCTGGCTGATCTGGG |
| Cavin4 | *Cavin4/ Murc* | 68016 | GTAATCTTCCAGGAGGACATTCC<br>CCGAGGAGAGCTCTATTGGG |
| Popeye domain containing 1 | *Popdc1/ Bves* | 23828 | GATGTTGTCTCTAGGATGTACCC<br>ATATTGATACCCAAGAACACCGAG |
| Popeye domain containing 2 | *Popdc2* | 64082 | TTTGAGGAGGTTTCAGGATCAG<br>GAGGACGTCTAGAGGGTAGG |
| Popeye domain containing 3 | *Popdc3* | 78977 | GCTCTGAAGTGGTTAGTTTGG<br>ACTGTCACTCTGATTCTTCCTG |
| MicroRNA predicted targeting Caveolin-3 | microRNA-22 | 387141 | AAGCTGCCAGTTGAAGA<br>Universal Primer |

| | | | |
|---|---|---|---|
| MicroRNA predicted targeting Caveolin-3 | microRNA-101b | 724062 | CAGTACTGTGATAGCTGAA |
| | | | Universal Primer |

**Supplementary Table 2. Detailed 2-Way ANOVA Output for Male Caveolin, Cavins and Popdcs.**

| Gene | Factor | Mean Square | F (DFn, DFd) | % of Total Variation | P value |
|---|---|---|---|---|---|
| ***Cav1*** | Age | 0.6608 | 5.56 (3, 40) | 25.6% | 0.0028 |
| | Ischemia | 0.06780 | 0.57 (1, 40) | 0.9% | 0.4547 |
| | Age*Ischemia | 0.3086 | 2.59 (3, 40) | 12.0% | 0.0658 |
| ***Cav2*** | Age | 0.3915 | 4.69 (3, 40) | 24.7% | 0.0067 |
| | Ischemia | 0.05610 | 0.67 (1, 40) | 1.2% | 0.4173 |
| | Age*Ischemia | 0.06175 | 0.74 (3, 40) | 3.9% | 0.5348 |
| ***Cav3*** | Age | 0.4257 | 13.33 (3, 40) | 27.4% | <0.0001 |
| | Ischemia | 1.050 | 32.87 (1, 40) | 22.5% | <0.0001 |
| | Age*Ischemia | 0.3516 | 11.01 (3, 40) | 22.6% | <0.0001 |
| ***Cavin1*** | Age | 1.014 | 11.03 (3, 40) | 43.2% | <0.0001 |
| | Ischemia | 0.1925 | 2.10 (1, 40) | 2.7% | 0.1555 |
| | Age*Ischemia | 0.04263 | 0.46 (3, 40) | 1.8% | 0.7090 |
| ***Cavin2*** | Age | 0.1761 | 5.25 (3, 40) | 14.4% | 0.0038 |
| | Ischemia | 1.184 | 35.30 (1, 40) | 32.3% | <0.0001 |
| | Age*Ischemia | 0.2020 | 6.02 (3, 40) | 16.6% | 0.0017 |
| ***Cavin3*** | Age | 0.1319 | 1.38 (3, 40) | 9.1% | 0.2626 |
| | Ischemia | 0.080 | 0.08 (1, 40) | 0.2% | 0.7740 |
| | Age*Ischemia | 0.0498 | 0.52 (3, 40) | 3.4% | 0.6699 |
| ***Cavin4*** | Age | 0.1202 | 2.46 (3, 38) | 12.8% | 0.0774 |
| | Ischemia | 0.0078 | 0.16 (1, 38) | 0.3% | 0.6914 |
| | Age*Ischemia | 0.1957 | 4.01 (3, 38) | 20.9% | 0.0142 |
| ***Popdc1*** | Age | 0.2113 | 4.88 (3, 37) | 18.5% | 0.0059 |
| | Ischemia | 0.6739 | 15.56 (1, 37) | 19.7% | 0.0003 |
| | Age*Ischemia | 0.1571 | 3.63 (3, 37) | 13.8% | 0.0216 |
| ***Popdc2*** | Age | 0.3636 | 1.21 (3, 37) | 8.4% | 0.3184 |
| | Ischemia | 0.1332 | 0.44 (1, 37) | 1.0% | 0.5091 |
| | Age*Ischemia | 0.2146 | 0.72 (3, 37) | 4.9% | 0.5486 |
| ***Popdc3*** | Age | 0.0277 | 0.19 (3, 35) | 0.8% | 0.8993 |
| | Ischemia | 3.725 | 26.20 (1, 35) | 37.6% | <0.0001 |
| | Age*Ischemia | 0.4659 | 3.28 (3, 35) | 14.1% | 0.0323 |

**Supplementary Table 3. Detailed 2-Way ANOVA Output for Female Caveolin, Cavins and Popdcs.**

| Gene | Factor | Mean Square | F (DFn, DFd) | % of Total Variation | P value |
|---|---|---|---|---|---|
| ***Cav1*** | Age | 1.210 | 4.36 (3, 38) | 20.0% | 0.0098 |
| | Ischemia | 0.8040 | 2.90 (1, 38) | 4.4% | 0.0967 |
| | Age*Ischemia | 0.9977 | 3.60 (3, 38) | 16.5% | 0.0220 |
| ***Cav2*** | Age | 0.8263 | 3.62 (3, 40) | 13.1% | 0.0211 |
| | Ischemia | 4.820 | 21.10 (1, 40) | 25.4% | <0.0001 |
| | Age*Ischemia | 0.8467 | 3.71 (3, 40) | 13.4% | 0.0192 |
| ***Cav3*** | Age | 1.397 | 6.91 (3, 38) | 19.7% | 0.0008 |
| | Ischemia | 8.158 | 40.43 (1, 38) | 38.3% | <0.0001 |
| | Age*Ischemia | 0.5714 | 2.82 (3, 38) | 8.1% | 0.0516 |
| ***Cavin1*** | Age | 0.3854 | 0.84 (3, 38) | 5.1% | 0.4815 |
| | Ischemia | 2.511 | 5.46 (1, 38) | 11.0% | 0.0248 |
| | Age*Ischemia | 0.5334 | 1.16 (3, 38) | 7.0% | 0.3377 |
| ***Cavin2*** | Age | 0.4823 | 4.00 (3, 40) | 16.5% | 0.0139 |
| | Ischemia | 0.0007 | 0.01 (1, 40) | 0.00% | 0.9400 |
| | Age*Ischemia | 0.8423 | 6.99 (3, 40) | 28.7% | 0.0007 |
| ***Cavin3*** | Age | 1.393 | 8.07 (3, 39) | 29.8% | 0.0003 |
| | Ischemia | 2.127 | 12.33 (1, 39) | 15.2% | 0.0011 |
| | Age*Ischemia | 0.3763 | 2.18 (3, 39) | 8.1% | 0.1058 |
| ***Cavin4*** | Age | 1.436 | 9.92 (3, 40) | 40.2% | <0.0001 |
| | Ischemia | 0.1895 | 1.31 (1, 40) | 1.8% | 0.2594 |
| | Age*Ischemia | 0.1411 | 0.97 (3, 40) | 4.0% | 0.4142 |
| ***Popdc1*** | Age | 1.335 | 3.63 (3, 40) | 14.4% | 0.0208 |
| | Ischemia | 2.307 | 6.28 (1, 40) | 8.3% | 0.0164 |
| | Age*Ischemia | 2.256 | 6.14 (3, 40) | 24.4% | 0.0016 |
| ***Popdc2*** | Age | 1.204 | 4.78 (3, 38) | 24.7% | 0.0063 |
| | Ischemia | 0.0917 | 0.36 (1, 38) | 0.6% | 0.5498 |
| | Age*Ischemia | 0.4646 | 1.85 (3, 38) | 9.5% | 0.1555 |
| ***Popdc3*** | Age | 0.8771 | 6.91 (3, 40) | 18.9% | 0.0007 |
| | Ischemia | 5.798 | 45.69 (1, 40) | 41.7% | <0.0001 |
| | Age*Ischemia | 0.1334 | 1.05 (3, 40) | 2.9% | 0.3806 |

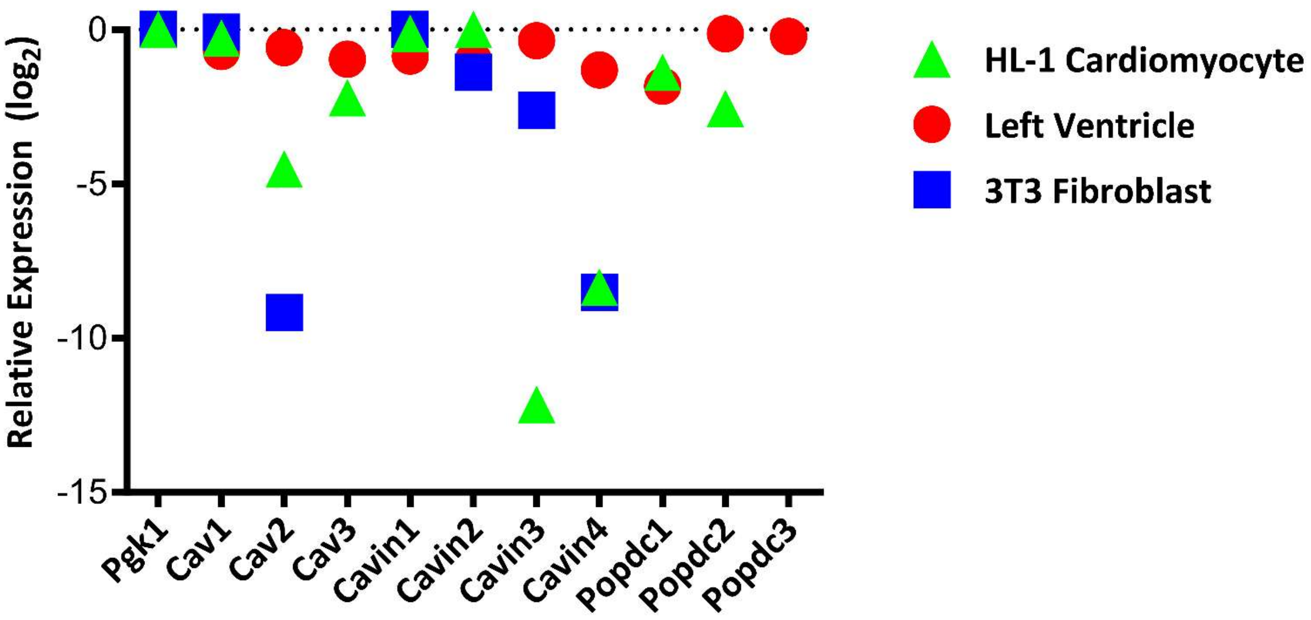


**Supplementary Figure S1. Relative expression of Caveolin, Cavin and Popdc transcripts in left ventricle, HL-1 cardiomyocytes and 3T3 fibroblasts**. The dot-plot presents relative caveolin, cavin and Popdc gene expression levels in left ventricular tissue (8-week old male mice), cultured HL-1 cardiomyocytes and cultured 3T3 fibroblasts. Data presented as means relative to *Pgk1* expression. (n=3/group).
